# Hippocampal γCaMKII dopaminylation promotes synaptic-to-nuclear signaling and memory formation

**DOI:** 10.1101/2024.09.19.613951

**Authors:** Andrew F. Stewart, Sasha L. Fulton, Romain Durand-de Cuttoli, Robert E. Thompson, Peng-Jen Chen, Elizabeth Brindley, Bulent Cetin, Lorna A. Farrelly, Rita Futamura, Sarah Claypool, Ryan M. Bastle, Giuseppina Di Salvo, Christopher Peralta, Henrik Molina, Erdene Baljinnyam, Samuele G. Marro, Scott J. Russo, Robert J. DeVita, Tom W. Muir, Ian Maze

**Author notes:** Corresponding authors, **Ian Maze, Email:****, Tom W. Muir, Email:**.

## Abstract

Protein monoaminylation is a class of posttranslational modification (PTM) that contributes to transcription, physiology and behavior. While recent analyses have focused on histones as critical substrates of monoaminylation, the broader repertoire of monoaminylated proteins in brain remains unclear. Here, we report the development/implementation of a chemical probe for the bioorthogonal labeling, enrichment and proteomics-based detection of dopaminylated proteins in brain. We identified 1,557 dopaminylated proteins – many synaptic – including γCaMKII, which mediates Ca^2+^-dependent cellular signaling and hippocampal-dependent memory. We found that γCaMKII dopaminylation is largely synaptic and mediates synaptic-to-nuclear signaling, neuronal gene expression and intrinsic excitability, and contextual memory. These results indicate a critical role for synaptic dopaminylation in adaptive brain plasticity, and may suggest roles for these phenomena in pathologies associated with altered monoaminergic signaling.

## MAIN TEXT

Biogenic amines, such as serotonin, dopamine and histamine, are a large class of evolutionarily ancient signaling molecules. These metabolites are found in diverse tissues in animals and can elicit biological effects in an auto-, para- or endocrine manner (*1, 2*). In brain, these monoamines are thought to function primarily via synaptic release, orchestrating diverse behaviors, including mood, appetite, pleasure, and sleep, as well as mediating cognitive functions, such as attention, learning, and memory (*3–5*). Altered monoaminergic signaling is linked to many mental health/neurological disorders, including depression and addiction, and in the case of dopamine, Parkinson’s disease (*6*). The canonical signaling pathways activated by biogenic amines are reasonably well understood (*7–9*) – they engage cognate cell surface receptors, typically G protein coupled receptors (GPCRs), of which there can be several members for each monoamine type, leading to activation of intracellular signaling cascades that elicit a number of cellular outputs (*10*). While these pathways rely on vesicular packaging and synaptic release of monoamines, it has long been recognized that non-vesicular pools of monoamines (both cytoplasmic and nuclear) exist in various cell-types, including neurons (*11, 12*). The existence of these pools suggested a non-canonical, receptor-independent role for monoamines in cell biology. Evidence for this goes back several decades, where it was found that biogenic amines (both mono- and polyamines) can be covalently attached to proteins by transglutaminase enzymes (*13–15*).

Motivated by these and other more recent findings (*16–18*), we previously asked whether this class of posttranslational modification (PTM) can occur on histones, the core packing proteins of chromatin. Tissue transglutaminase 2 (TGM2)-mediated serotonylation (*19–24*), dopaminylation (*25–27*) and histaminylation (*28*) of histone H3 have all now been documented and are found in both monoaminergic and non-monoaminergic cells of the brain. The genomic enrichment patterns and overall dynamics of these histone monoaminylations argue for critical roles in transcriptional, physiological and behavioral plasticity as part of both normal neural development, and in the context of brain pathologies. However, given the enzymatic promiscuity of TGM2, it seemed possible – perhaps even likely based upon our previous data – that glutamine-containing proteins in other cellular compartments in brain may also be subject to monoaminylation. However, the full extent of protein monoaminylations across neural tissues remains unknown. By extension, it is also unclear how the levels or prevalence of these modifications may change as a function of cell/tissue state, whether that be through differentiation, activation of signaling pathways or environmental perturbations. As such, in this study we sought to develop an approach that would allow for the identification of the broader repertoire of monoaminylated proteins in brain (focusing here on protein dopaminylation) to more fully understand the contributions of these non-canonical monoamine signaling moieties in brain health and disease.

### Design of a chemical probe for the bioorthogonal labeling, enrichment and detection of dopaminylated proteins

Inspired by earlier work (*29, 30*) – and in agreement with recent *in vitro* findings (*31, 32*) – demonstrating that 1,2 catechols (e.g., dopamine/DA, norepinephrine/NE) can undergo a strain-promoted cycloaddition reaction with cyclooctynes following oxidation to orthoquinones, we developed a modified bioorthogonal strategy whereby periodate treated catecholaminylated peptides can be efficiently labeled with a biotinylated strained cyclooctyne probe (Bio-CO) (**Fig. S2A**); see **Fig. S1A-C** for synthesis characterization. First, to validate that a known substrate of protein dopaminylation can be efficiently IP’d from neural tissues, Bio-CO labeling and enrichment (using Streptavidin as prey) of histone H3 from mouse brain extracts was performed, which confirmed that catecholaminylated H3 was robustly labeled by the probe (+ probe *vs.* – probe comparisons are provided to demonstrate a lack of background signal in the absence of probe labeling) (**Fig. S2B**). To then assess the specificity of our probe against different monoaminylated substrates, we performed *in vitro* transglutaminase reactions against recombinant histone H3 using various monoamine donors (serotonin/5-HT, histamine/His, DA and NE), followed by Bio-CO enrichment and western blotting for H3. Importantly, since H3 can be modified by multiple catecholamines (DA and NE; see **Fig. 2C-F** for *in vitro* validations of H3Q5 noradrenylation) in a TGM2-dependent manner, we wished to explore whether the Bio-CO probe equally labels a dopaminylated *vs.* noradrenylated substrate. We found that Bio-CO efficiently immunoprecipitated (IP’d) dopaminylated H3, with no labeling observed for other monoaminylated forms of the protein, including noradrenylated H3 (**Fig. S2G**). While, in theory, Bio-CO would also be predicted to label noradrenylated substrates, we found that under the specific reaction conditions used in this study that no such labeling was observed, possibly owing to steric hindrance between the probe and the β-hydroxyl of NE (**Fig. S2H**), although the precise reasons for this lack of labeling remain unclear. While further optimization of reaction conditions may result in efficient labeling of noradrenylated proteins, we viewed such selectivity towards dopaminylated substrates as an advantage in our study. To further assess the efficacy and selectivity of the Bio-CO probe in labeling dopaminylated *vs.* noradrenylated proteins in neurons, we treated cultured primary (DIV14) mouse corticostriatal neurons (which, *in vivo*, do not synthesize the monoamines but rather receive projections from catecholaminergic neurons) with 500 nM DA or NE for 1 hr, followed by Bio-CO labeling, Streptavidin enrichment and mass spectrometry (LC-MS/MS) (**Fig. S2I**). Similar to our *in vitro* findings, these results indicated that Bio-CO robustly labeled dopaminylated (**Fig. S2J, Data S1)**, but not noradrenylated (**Fig. S2K, Data S2)**, proteins in treated neurons. 330 uniquely dopaminylated proteins were identified by LC-MS/MS in this culture system, the vast majority of which were found to be synaptic (both pre- and post-synaptic; SynGO 2024) and/or involved in synaptic regulation (GO BP 2023), and were enriched in biological pathways known to be disrupted in brain disorders (DisGeNET) associated with altered dopamine signaling – e.g., schizophrenia, intellectual disability, Parkinson’s disease, etc. (**Fig. S2L**).

### Characterization of the mouse brain dopaminylome *in vivo*

Given the relative immaturity of neuronal primary culture systems, as well as the relative absence of other relevant neural cell-types (e.g., glia), we next sought to characterize the *in vivo* protein dopaminylome in mouse brain. Adult tissues from mouse ventral tegmental area (VTA), nucleus accumbens (NAc), medial prefrontal cortex (mPFC) and dorsal hippocampus (dHPC) were chosen as brain regions of interest owing to their known DA-dependent physiology and their relevance to DA-mediated behaviors (*33*). Following Bio-CO labeling of mouse brain extracts, Streptavidin enrichment and LC-MS/MS, we identified a total of 1,557 uniquely dopaminylated substrates across these four brain regions, with VTA – which is enriched for dopaminergic neurons – displaying the largest number of dopaminylated proteins (1,281; **Fig. 1A, Data S3**), followed by NAc (737; **Fig. 1B, Data S4**), mPFC (719; **Fig. 1C, Data S5**) and dHPC (357; **Fig. 1D, Data S6**), the latter three of which receive dense (NAc, mPFC) or restricted (dHPC) (*34*) DA projections from VTA (as well as substantia nigra, which was not investigated here). Similar to our *in cellulo* results using primary cultured neurons, a large percentage of these substrates were found to be synaptic (both pre- and post-synaptic; SynGO 2024), involved in synaptic regulation (GO BP 2023) and were enriched in biological pathways known to be disrupted in brain disorders (DisGeNET) that are associated with altered dopamine signaling – e.g., schizophrenia, intellectual disability, Parkinson’s disease, etc. (**Fig. 1E**). While some of the dopaminylated proteins were found to be uniquely modified in a brain region-specific manner, many of the proteins were found to be commonly modified across brain structures, with 91 putatively modified substrates observed in all four brain regions (**Fig. 1F**). Among these 91 proteins were numerous important synaptic constituents/regulators, including Dlg4 (PSD-95), NMDA receptor subunits Grin1 and Grin2b, and three members of the Ca^2+^/calmodulin-dependent protein kinase family – αCaMKII (Camk2a), βCaMKII (Camk2b) and γCaMKII (Camk2g) (∂CaMKII, while expressed in brain, was only shown to be a substrate of dopaminylation in VTA), the latter three of which were validated in mouse brain extracts (dHPC) via Bio-CO labeling, enrichment and western blotting (**Fig. 1G**). To further confirm that our Bio-CO probe labeled endogenously dopaminylated proteins and not those that may be modified *in situ* following tissue/cell lysis (during which time TGM2 and DA may theoretically come into contact with proteins with which they may otherwise not interact *in vivo*), we performed a spike-in control experiment using recombinant, non-dopaminylated FLAG-tagged γCaMKII, which was added to tissue extracts post-lysis, followed by Bio-CO labeling, enrichment and western blotting for FLAG or γCaMKII. These results confirmed that the endogenous γCaMKII dopaminylation observed in **Fig. 1A-D, G** was not due to TGM2-mediated dopaminylation *in situ* (**Fig. S2M**). Next, to ensure that the dopaminylation observed for αCaMKII, βCaMKII and γCaMKII was direct (i.e., the proteins were not IP’d as a result of being bound to other dopaminylated proteins in the extracts), we performed *in vitro* TGM2 enzymatic assays on recombinant αCaMKII, βCaMKII and γCaMKII using monodansylcadaverine (MDC; an autofluorescent monoamine analog) with or without donor competition using excess DA. These results indicated that all three CaMKII proteins were direct substrates of TGM-dependent dopaminylation (**Fig. 1H**).

**Fig. 1:**
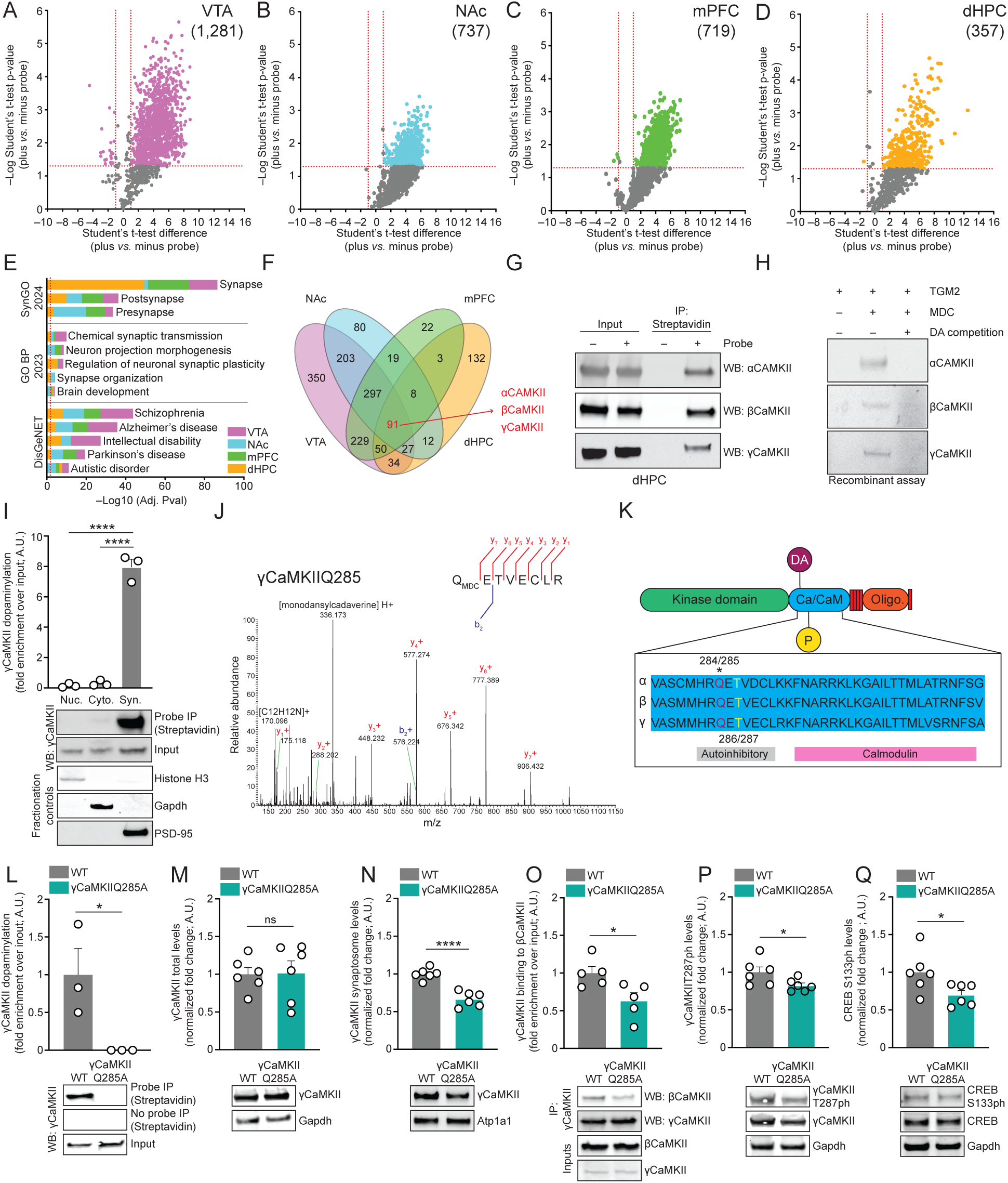
Global proteomic profiling in mammalian brain identifies roles for protein dopaminylation in synaptic-to-nuclear signaling. Volcano plots of MS-identified dopaminylated proteins enriched by the Bio-CO probe in (**A**) VTA, (**B**) NAc, (**C**) mPFC and (**D**) dHPC (Student’s t-test corrected for multiple comparisons; + probe *vs.* – probe: *n* = 3; *p*<0.05, fold-change >1). (**E**) Ontology analysis (SynGO 2024, GO BP 2023, DisGeNET) of MS-identified proteins enriched by the Bio-CO probe in VTA, NAc, mPFC and dHPC (FDR<0.05; Benjamini-Hochberg). (**F**) Venn diagram depicting the overlap of MS-identified dopaminylated proteins enriched by the Bio-CO probe in VTA *vs.* NAc *vs.* mPFC *vs.*dHPC (from **A**). (**G**) Bio-CO probe-mediated bioorthogonal labeling of endogenously dopaminylated αCaMKII, βCaMKII and γCaMKII from mouse dHPC. Inputs = 1%; Streptavidin IPs were performed for – *vs.* + probe conditions, followed by blotting for αCaMKII, βCaMKII and γCaMKII. (**H**) Bio-CO probe-mediated bioorthogonal labeling of MDCylated αCaMKII, βCaMKII and γCaMKII *in vitro* following TGM2-dependent transglutamination. DA competition revealed that MDCylation of recombinant αCaMKII, βCaMKII and γCaMKII effectively eliminates MDCylation signal for all three proteins. (**I**) Bio-CO probe-mediated bioorthogonal labeling of endogenously dopaminylated γCaMKII from sub-cellular fractions of dHPC revealed that γCaMKII dopaminylation is exclusively synaptic. Fractionation control blots for the sub-cellular compartments (H3 for nuclear, Gapdh for cytosolic and PSD-95 for synaptic) are provided, and γCaMKII levels were normalized to respective inputs (1%). *n* = 3/fraction – significance determined by one-way ANOVA (F_2,6_ = 215.9, \*\*\*\**p*<0.0001), followed by post hoc analysis (Tukey’s MC test, \*\*\*\**p*<0.0001). (**J**) MS spectra for MDCylation of γCaMKII at glutamine 285. Y+ and b+ ions are annotated in red and blue, respectively. (**K**) Cartoon of protein sequence alignment and domain structure for αCaMKII *vs.* βCaMKII *vs.* γCaMKII. (**L**) Bio-CO probe-mediated bioorthogonal labeling of dopaminylated γCaMKII from dHPC of wildtype *vs.* γCaMKIIQ285A mutant mice. Streptavidin IPs were performed for – *vs.* + probe conditions and γCaMKII levels were normalized to respective inputs (1%). *n* = 3/genotype – significance determined by unpaired Student’s t-test (t_4_ = 2.873, \**p* = 0.0453). (**M**) Western blotting for total γCaMKII levels (normalized to Gapdh as a loading control) in dHPC from wildtype *vs.* γCaMKIIQ285A mutant mice. *n* = 6/genotype – significance determined by unpaired Student’s t-test (t_10_ = 0.1185, *p* = 0.9080). (**N**) Western blotting for total γCaMKII levels in dHPC synaptic fractions (normalized to Atp1a1 as a loading control) from wildtype *vs.* γCaMKIIQ285A mutant mice. *n* = 6/genotype – significance determined by unpaired Student’s t-test (t_10_ = 6.592, \*\*\*\**p*<0.0001). (**O**) Western blotting of dHPC co-IPs to assess βCaMKII binding to γCaMKII in wildtype *vs.* γCaMKIIQ285A mutant mice. *n* = 5/genotype – significance determined by unpaired Student’s t-test (t_8_ = 2.711, \**p* = 0.0266). Levels of βCaMKII binding to γCaMKII were normalized to respective inputs (1%). (**P**) Western blotting for γCaMKIIT287ph levels (normalized to total γCaMKII levels and Gapdh as loading controls) in dHPC from wildtype *vs.* γCaMKIIQ285A mutant mice. *n* = 6/genotype – significance determined by unpaired Student’s t-test (t_10_ = 2.240, \**p* = 0.0490). (**Q**) Western blotting for CREBS133ph levels (normalized to total CREB levels and Gapdh as loading controls) in dHPC from wildtype *vs.* γCaMKIIQ285A mutant mice. *n* = 6/genotype – significance determined by unpaired Student’s t-test (t_10_ = 2.361, \**p* = 0.0399). All bar plots presented as mean ± SEM. See **Figure S8** for uncropped blots.

**Fig. 2:**
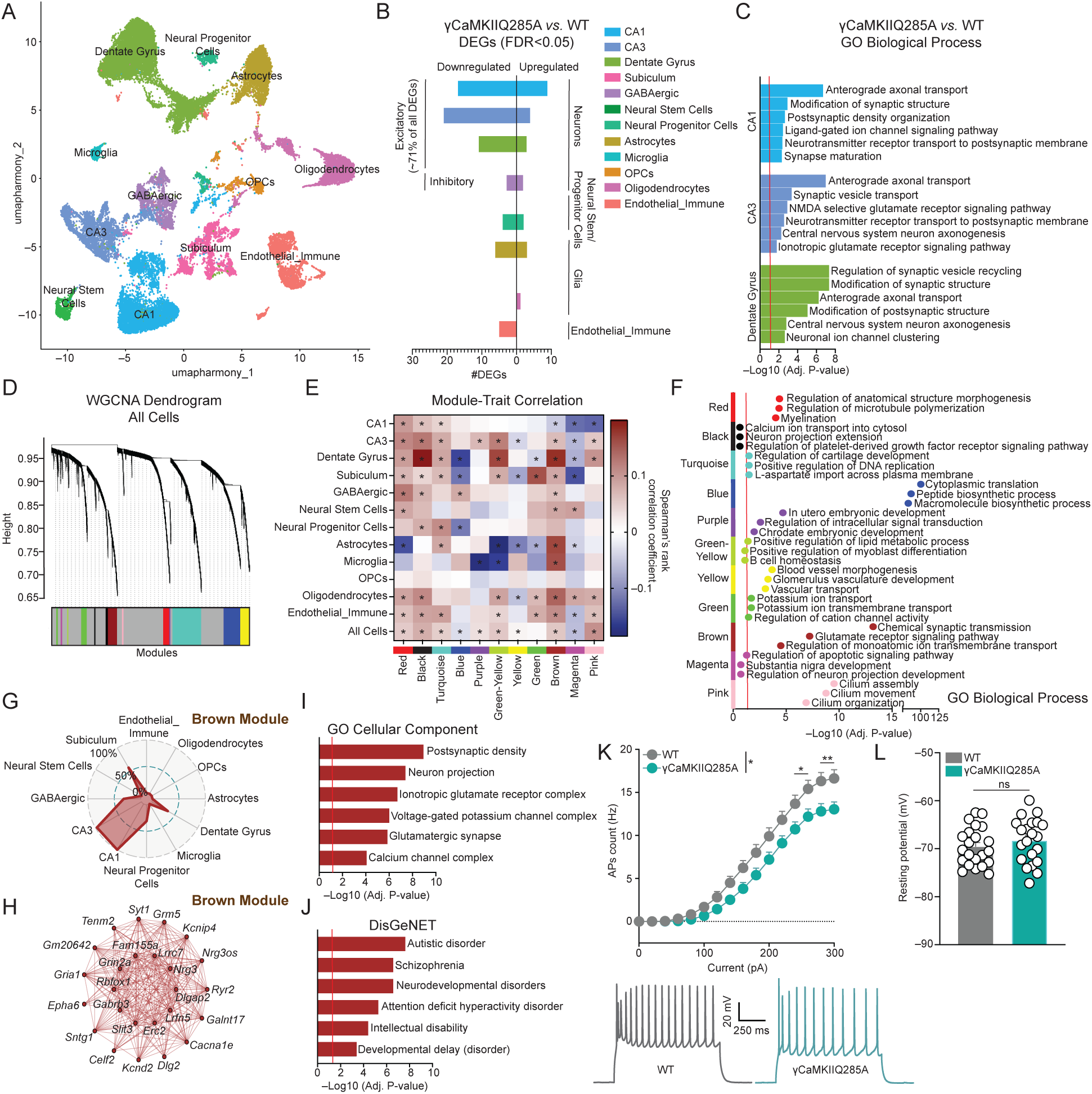
Synaptic γCaMKIIQ285dop is required for normal hippocampal gene expression and intrinsic CA1 pyramidal neuron excitability. (**A**) Uniform manifold approximation and projection (UMAP), colored by major cell-type (*n* = 3 wildtype; *n* = 4 γCaMKIIQ285A). (**B**) Number of DEGs observed by cell-type from pseudobulk analysis (FDR<0.05). (**C**) Gene set enrichment analysis (GO Biological Process) for DEGs identified in CA1, CA3 and Dentate Gyrus excitatory neurons from pseudobulk analysis (FDR<0.05; Benjamini-Hochberg). (**D**) Dendrogram showing co-expression modules identified by weighted gene correlation network analysis (WGCNA). (**E**) Heatmap of co-expression module correlation with mutation (γCaMKIIQ285A) trait by cell-type. * indicates Adj. P <. 05 significance of correlation. (**F**) enrichR-based gene ontology (GO Biological Process) analysis for genes in significant co-expression modules (FDR<0.05; Benjamini-Hochberg). (**G**) Radar plots of WGCNA-identified Brown module expression by annotated cell cluster. % expression of module genes by cell-type is noted. (**H**) Network underlying the top 25 hub genes in the Brown module. Each node represents a hub gene, and each edge represents the co-expression relationship between two genes in the network. The top 10 hub genes are in the center of the plot, while the remaining 15 are in the outer circle. (**I**) enrichR-based gene ontology (GO Cellular Component) analysis for genes in brown module (FDR<0.05; Benjamini-Hochberg). (**J**) enrichR-based gene ontology (DisGeNET) analysis for genes in brown module (FDR<0.05; Benjamini-Hochberg). (**K**) The number of evoked action potentials (APs) in dHPC CA1 pyramidal neurons of wildtype (*n* = 22 neurons from 5 mice) *vs.* γCaMKIIQ285A (*n* = 21 neurons from 4 mice) in response to increasing depolarizing current steps. Representative membrane responses are provided from dHPC CA1 pyramidal neurons in response to a 280 pA current injection in wildtype or γCaMKIIQ285A mutant mice. Significance was determined via two-way ANOVA (main effect of interaction between current injection and genotype; F_15, 656_ = 1.814) with subsequent post hoc analysis (Sidak’s MC test: \**p* = 0.0425 or 0.0182, \*\**p* = 0.0064 or 0.0047). (**L**) Resting membrane potential of dHPC CA1 pyramidal neurons of wildtype (*n* = 22 neurons from 5 mice) *vs.* γCaMKIIQ285A (*n* = 21 neurons from 4 mice). Significance was determined by unpaired Student’s t-test (t_41_ = 0.9639, *p* = 0.3407). All bar/line plots presented as mean ± SEM.

Given that our proteomics data robustly implicated roles for brain dopaminylation in synaptic development, regulation and plasticity, along with their relative enrichment in disease ontologies related to cognition (e.g., intellectual disability, autism spectrum disorder, etc.), we next wished to further dissect potential mechanistic roles for these dopaminylation events *in vivo*. One particular validated candidate of note, γCaMKII, had recently been shown to be member of a cluster of genes (*CAMK2G*) associated with memory in humans and was shown to at least partially explain variation in human hippocampal activity via fMRI (*35*). Furthermore, alternative splicing events in *CAMK2G* have been shown to be associated with autism (*36*), and γCaMKIIR292P mutations are known to result in profound synaptic deficits and intellectual disability in humans, with expression of γCaMKIIR292P in mice similarly yielding memory deficits (*37, 38*). Finally, in exhaustive work from the Tsien lab, γCaMKII was previously demonstrated to be a critical synaptic-to-nuclear signaling molecule, where it translocates from the cytosol to the cell membrane upon Ca^2+^-dependent activation of βCaMKII, resulting in its further shuttling with calmodulin (CaM) to the nucleus to phosphorylate the transcription factor CREB. These events, in turn, induce transcription at CREB target genes (*39*). Importantly, Ma *et al.,* demonstrated that γCaMKII shuttling to the cell membrane, which allows for direct interactions with βCaMKII, functions as a critical ‘checkpoint’ for signaling to proceed properly (*39*). Despite this incredibly thorough work, however, the signal for recruitment of γCaMKII to the synaptic membrane upon cellular activation was not identified. Given that γCaMKII can exist in the cytoplasm, the nucleus and at the synapse, we next aimed to examine in which sub-cellular compartment γCaMKII is dopaminylated – focusing now on the hippocampus given γCaMKII’s important regulatory roles in this region – to assess whether this modification may influence γCaMKII’s cellular signaling, events that are critical for hippocampal-dependent learning and memory. As such, we biochemically fractionated dHPC tissues from wildtype adult mice into nuclear-, cytosolic- and synaptosomal-enriched fractions, followed by Bio-CO labeling, enrichment and western blotting for γCaMKII. Our results indicated that γCaMKII dopaminylation was almost exclusively synaptic (**Fig. 1I**). We next explored which amino acid residue(s) within γCaMKII are direct substrates for monoaminylation. We again performed TGM2 enzymatic assays against recombinant human γCaMKII using MDC as a monoamine donor, followed by LC-MS/MS. While γCaMKII contains numerous glutamines (Q) (the primary amino acid substrate for transglutamination), we found that only one residue – γCaMKIIQ285 – was able to be modified *in vitro* (**Fig. 1J**). To confirm whether the specificity for Q285 monoaminylation was consistent across other CaMKII proteins, we similarly performed TGM2 enzymatic assays against αCaMKII, where we found that this same glutamine (Q284 in αCaMKII) was the primary site of monoaminylation (**Fig. S3A-B**). Since γCaMKIIQ285 exists within a conserved region of the CaMKII kinase family that functions as a Ca^2+^-CaM binding domain (**Fig. 1K**), we reasoned that dopaminylation of this glutamine may impact aspects of its cellular signaling.

### Synaptic γCaMKIIQ285dop contributes to γCaMKII membrane localization and downstream synaptic-to-nuclear signaling

As alluded to previously, γCaMKII is a synaptic-to-nuclear signaling protein that sequesters CaM in a threonine 287 phosphorylation (T287ph)-dependent manner, thereby promoting nuclear shuttling, CREB phosphorylation at serine 133 (ser133ph) and CREB-mediated transcription. Given that γCaMKIIQ285 is only two amino acid residues away from the autoinhibitory T287ph site, we hypothesized that γCaMKIIQ285dop may promote crosstalk interactions between these two PTMs to regulate γCaMKII signaling through this pathway. To further investigate this potential crosstalk mechanism *in vivo*, we next employed CRISPR-based gene editing in mice to generate whole-body γCaMKIIQ285A – a mutation that does not impact γCaMKII’s intrinsic kinase activity *in* vitro; (**Fig. S3C**) – knock-in mice (**Fig. S3D-G**), which completely abolished endogenous γCaMKIIQ285dop levels in mutant animals (**Fig. 1L**). Next, to assess what impact eliminating γCaMKIIQ285dop may have on γCaMKII signaling in brain, we performed a series of biochemical assessments on dHPC tissues from wildtype *vs.* γCaMKIIQ285A mice, which demonstrated that while the mutation does not impact total γCaMKII levels (**Fig. 1M**; nor total levels of protein dopaminylation in dHPC – **Fig. S3H**), loss of the mark resulted in reduced trafficking of γCaMKII to the synapse (**Fig. 1N**), decreased binding of γCaMKII to βCaMKII (**Fig. 1O**), attenuation of γCaMKIIT287ph (**Fig. 1P**) and reduced levels CREBS133ph (**Fig. 1Q**). To further validate the potential impact of reducing γCaMKIIQ285dop levels on CREB-mediated transcription, we transduced primary hippocampal neurons isolated from wildtype *vs.* γCaMKIIQ285A mice with a lentivirus expressing a CREB response element (CRE) driving a luciferase reporter gene, and then measured CREB-mediated transcription directly following treatment with 500 nM DA (*vs.* vehicle). While DA treatments in wildtype neurons resulted in a predictable increase in CREB-induced transcription, we observed a significant attenuation of this activity in γCaMKIIQ285A neurons (**Fig. S3I**). These results indicated that γCaMKIIQ285dop contributes importantly to γCaMKII’s synaptic-to-nuclear signaling and downstream CREB-mediated transcription, phenomena that may further impact neuronal gene expression, physiology and/or behavior.

### γCaMKIIQ285dop influences neuronal gene expression in hippocampus and contributes to CA1 neuron excitability

We next wished to examine how reducing levels of the mark affects gene expression across hippocampal cell-types. For this, we performed single-nuclei RNA-sequencing (snRNA-seq) on adult dHPC tissues from wildtype *vs.* γCaMKIIQ285A mutant mice. After removing doublets and filtering out low quality cells (see **Fig. S4A-D** for relevant quality control metrics post-filtering), we obtained a total of 33,983 nuclei for downstream processing (wildtype = 16,256; γCaMKIIQ285A = 17,727). Using both manual curation and label transfer with high-resolution validated annotations from previously published hippocampus datasets from the Allen Brain Atlas (*40*), we annotated 12 major cell-type groups, including excitatory neuron subtypes (CA1, CA3, Dentate Gyrus and Subiculum; 20,510 nuclei, 60.35% of total), GABAergic subtypes (2,040 nuclei, 6.00% of total), neural stem cells and neural progenitor cells (1,562 nuclei, 4.59% of total), glial cell-types (astrocytes, microglia, oligodendrocyte progenitor cells/OPCs and oligodendrocytes; 7,384 nuclei, 21.72% of total) and endothelial/immune cells (2,487 nuclei, 7.31% of total) (**Fig. 2A**, **Fig. S5A-C**). We next used pseudobulk aggregation and differential expression testing to identify differentially expressed genes (DEGs; FDR<0.05) between wildtype *vs.* γCaMKIIQ285A mutant mice within each annotated cell-type. Consistent with γCaMKII being broadly expressed across all cell-types (**Fig. S6A**), this analysis showed that while many cell-clusters displayed at least modest levels of differential gene expression as a result of reducing γCaMKIIQ285dop levels, excitatory neuron clusters (specifically CA1, CA3 and Dentate Gyrus) exhibited the most robust alterations (∼71% of all DEGs across all cell-types), with the vast majority of these genes being downregulated in mutant animals (**Fig. 2B, Data S7**); note that *Camk2g* expression itself was not found to be affected by the mutation, which is consistent with our western blotting results in **Fig. 1**. Gene set enrichment analysis with GO Biological Process datasets further demonstrated that gene sets related to synaptic function (e.g., anterograde axonal transport, synaptic vesicle transport, ionotropic glutamate receptor signaling pathway, ligand-gated ion channel signaling pathway, etc.) were significantly differentially enriched in CA1, CA3 and Dentate Gyrus between the two conditions (**Fig. 2C**).

Given the potential complexity and interconnectedness of transcriptional changes that may occur as a result of reducing γCaMKII dopaminylation, we next sought to examine the effects of the γCaMKIIQ285A mutation on network-level gene expression, rather than individual gene features. Therefore, we employed weighted gene correlation network analysis on our snRNA-seq dataset (hdWGCNA package; version 0.3.03) to identify cell-type specific modules of highly co-expressed genes across both conditions (**Fig. 2D**). We next evaluated if these identified modules were relevant to the γCaMKIIQ285A mutation by calculating the correlation between the expression module genes and the trait of mutation across identified cell-types (**Fig. 2E**). We found that several modules were significantly correlated with γCaMKIIQ285A, and that the directionality of module gene expression was cell-type specific. For example, the Brown module was positively correlated with the mutation in all cell-types (including CA3 and Dentate Gyrus) except for CA1 cells, where genes in this module showed a negative correlation with the mutation (corresponding to reduced expression in the γCaMKIIQ285A group). To better understand the functional relevance of these significantly correlated modules in excitatory neuron sub-types, we next performed gene ontology enrichment analysis (**Fig. 2F**). Of particular interest, the Brown module was enriched for genes related to synaptic function (e.g., chemical synaptic transmission, glutamate receptor signaling pathway, regulation of monoatomic ion transmembrane transport, etc.). Furthermore, genes in the Brown module were mainly expressed in excitatory neurons (CA1 and CA3 – **Fig. 2G-H**; see **Fig. S6B** for module enrichment across other annotated cell-types, and **Data S8** for lists of genes enriched in each module). These data suggested that genes within the Brown module may be uniquely disrupted in CA1 excitatory neurons by loss of γCaMKIIQ285dop and may, in turn, contribute to synaptic dysregulation in these neurons. To examine this possibility further and to pinpoint specific cellular components and disease ontologies related to these disrupted genes in CA1, we performed an expanded GO analysis on genes in the Brown module focusing on cellular localization/components (GO Cellular Component) and disease ontologies (DisGeNET). These analyses revealed significant correlations with synaptic structures, such as ionotropic glutamate receptor complexes, voltage-gated K^+^ channel complexes and Ca^2+^-channel complexes (**Fig. 2I**), as well as disease ontologies related to cognitive impairment (e.g., intellectual disability, autistic disorder, developmental delay; **Fig. 2J**).

In total, these sequencing data indicated significant deficits in dHPC excitatory neuron gene expression in mutant mice, as well as unique patterns of transcriptional dysregulation in CA1 that may be predicted to result in abnormal synaptic architecture (e.g., postsynaptic density) and/or disruptions in voltage-gated potassium/ionotropic glutamate receptor signaling. Given this, as well as previous data implicating γCaMKII in human hippocampal activity^33^, we next wished to explore what impact disruptions of the mark may have on intrinsic electrophysiological properties of CA1 neurons. For this, we performed *ex vivo* patch-clamp recordings on CA1 pyramidal neurons in wildtype *vs.* γCaMKIIQ285A mice to assess the impact of γCaMKIIQ285dop loss on intrinsic neuronal excitability. As predicted, these results indicated that dHPC CA1 intrinsic neuronal excitability was significantly attenuated in mutant animals (**Fig. 2K**; note that the resting membrane potential of CA1 neurons was not impacted by the mutation – **Fig. 2L**). Taken together, these data demonstrated that the transcriptional deficits induced by loss of γCaMKIIQ285dop-mediated synaptic-to-nuclear signaling directly impacted normal cellular physiology, which, in turn, may also be predicted to result in deficits of other dHPC-dependent phenotypes (i.e., behavior).

### γCaMKIIQ285dop in dHPC is critical for contextual memory formation

Given roles for γCaMKIIQ285dop in mediating γCaMKII trafficking to the synapse (and regulation of downstream signaling), hippocampal transcription related to synaptic function and intellect, and normal neuronal physiology, we next sought to explore what impact disrupting γCaMKIIQ285dop may have on cognition. We first asked whether levels of γCaMKIIQ285dop may be altered in response to a learned experience by putting wildtype mice through contextual fear conditioning. In this paradigm, animals were first habituated to a neutral environment before receiving a series of five intermittent shocks at random intervals (two seconds each, 0.9 mA) over the course of the training session (**Fig. 3A**). One hour following the final shock, dHPC tissues from trained (context + shock) *vs.* untrained (context only) animals were collected and subjected to Bio-CO labeling, enrichment and western blotting for γCaMKII (**Fig. 3B**) or βCaMKII (**Fig. S7A**). These results demonstrated that γCaMKIIQ285dop, but not βCaMKII dopaminylation, levels were induced in dHPC by context + shock training, suggesting that learning-induced increases in γCaMKIIQ285dop may contribute to cellular signaling/transcription that might be necessary for contextual learning and/or formation of contextual memories. To explore this further, wildtype *vs*.

**Fig. 3:**
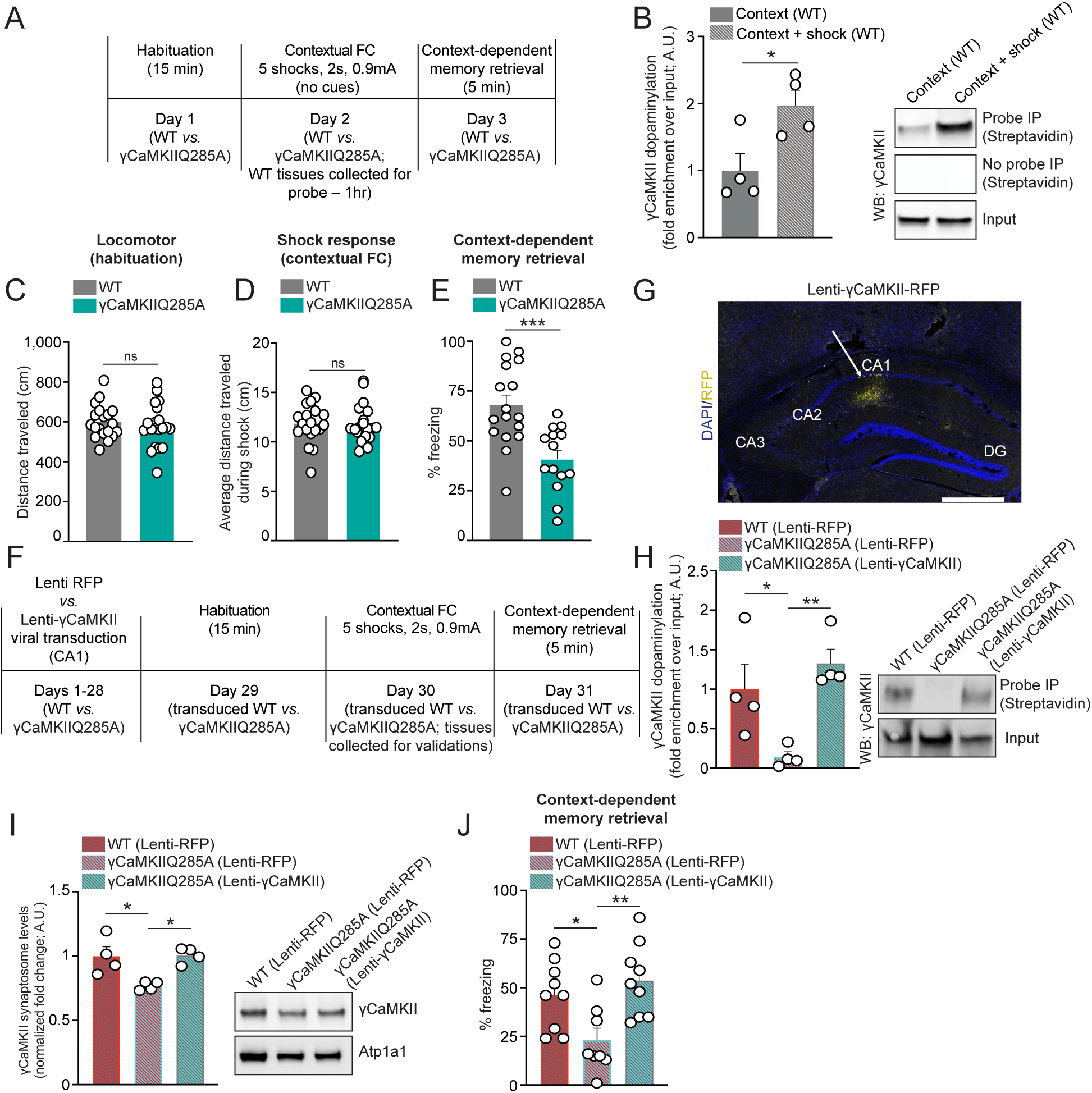
γCaMKIIQ285dop is induced during learning and contributes to contextual memory. (**A**) Timeline of contextual fear conditioning experiments in wildtype *vs.* γCaMKIIQ285A mutant mice. (**B**) Bio-CO probe-mediated bioorthogonal labeling of dopaminylated γCaMKII from dHPC of wildtype mice + (context + shock) *vs.* – (context alone) contextual fear conditioning training (1 hr post-training). Streptavidin IPs were performed for – *vs.* + probe conditions and γCaMKII levels were normalized to respective inputs (1%). *n* = 4/genotype – significance determined by unpaired Student’s t-test (t_6_ = 2.845, \**p* = 0.0294). (**C**) Locomotor activity during context habituation pre-fear conditioning for wildtype (*n* = 19) *vs.* γCaMKIIQ285A mutant (*n* = 20) mice. Significance determined by unpaired Student’s t-test (t_37_ = 0.6553, *p* = 0.5163). (**D**) Average locomotor activity across shock pairing trials for wildtype (*n* = 19) *vs.* γCaMKIIQ285A mutant (*n* = 20) mice. Significance determined by unpaired Student’s t-test (t_37_ = 0.1853, *p* = 0.8540). (**E**) Context-dependent memory retrieval 24 hr post-training for wildtype (*n* = 17) *vs.* γCaMKIIQ285A mutant (*n* = 14) mice. Significance determined by unpaired Student’s t-test (t_29_ = 4.002, \*\*\**p* = 0.0004). (**F**) Timeline of contextual fear conditioning experiments in wildtype *vs.* γCaMKIIQ285A mutant mice following intra-dHPC transduction with either Lenti-RFP or Lenti-γCaMKII. (**G**) IHC/IF image of Lenti-γCaMKII-RFP targeting to dHPC (CA1). DAPI was used as a counterstain to mark nuclei. Scale bar is set to 500 µm. (**H**) Bio-CO probe-mediated bioorthogonal labeling of dopaminylated γCaMKII from dHPC of wildtype *vs.* γCaMKIIQ285A mutant mice transduced with either Lenti-RFP or Lenti-γCaMKII. γCaMKII levels were normalized to respective inputs (1%). *n* = 4/genotype – significance determined by one-way ANOVA (F_2,9_ = 8.207, \*\**p* = 0.0094), followed by post hoc analysis (Tukey’s MC test; \**p* = 0.0468, \*\**p* = 0.0088). (**I**) Western blotting for total γCaMKII levels in dHPC synaptic fractions (normalized to Atp1a1 as a loading control) from wildtype *vs.* γCaMKIIQ285A mutant mice transduced with either Lenti-RFP or Lenti-γCaMKII. *n* = 4/genotype – significance determined by one-way ANOVA (F_2,9_ = 8.449, \*\**p* = 0.0086), followed by post hoc analysis (Tukey’s MC test; \**p* = 0.0163 for wildtype RFP *vs.* Q285A RFP, \**p* = 0.0143 for Q285A RFP *vs.* Q285A γCaMKII). (**J**) Context-dependent memory retrieval 24 hr post-training for wildtype (*n* = 9; RFP) *vs.* γCaMKIIQ285A mutant mice transduced with either Lenti-RFP (*n* = 8) or Lenti-γCaMKII (*n* = 9). Significance determined by one-way ANOVA (F_2,23_ = 6.676, \*\**p* = 0.0052), followed by post hoc analysis (Tukey’s MC test; \**p* = 0.0350, \*\**p* = 0.0049). All bar plots presented as mean ± SEM. See **Figure S8** for uncropped blots.

γCaMKIIQ285A mice were put through the same contextual fear conditioning paradigm and behavioral responses (locomotor behavior during habituation, shock response, learning acquisition and context-dependent memory retrieval) were assessed. Consistent with our molecular and biochemical data, we found that while γCaMKIIQ285A animals did not display alterations in locomotor behavior during habituation (**Fig. 3C**) or directly following shocks (**Fig. 3D**), they did display deficits in both learning acquisition [**Fig. S7B;** note that mutant mice did learn (reaching a threshold of ∼60% freezing by trial five), yet at a slower rate] and context-dependent memory retrieval (**Fig. 3E**).

Although these data clearly indicated that γCaMKIIQ285dop plays a critical role in contextual memory formation, it was important to consider that the γCaMKIIQ285A knock-in mice carry this mutation throughout their entire body, making it difficult to ascertain whether such memory deficits are indeed precipitated directly through alterations in hippocampal function. As such, we next wished to examine whether rescue of γCaMKIIQ285dop specifically within dHPC (focusing on CA1, a brain region/neuronal cell cluster most robustly impact by loss of the mark – see **Fig. 2**) could restore normal contextual memory in mutant mice. To do so, we transduced γCaMKIIQ285A mutant mice with a lentivirus expressing wildtype γCaMKII (co-expressing the fluorescent reporter RFP; Lenti-γCaMKII) *vs.* a lentivirus expressing RFP alone (Lenti-RFP), followed by 28 days of incubation to allow for maximal viral expression and then exposure to contextual fear conditioning. A separate group of wildtype mice were transduced with Lenti-RFP to serve as an additional control in our experimental design (**Fig. 3F**). In an independent cohort of animals not exposed to contextual fear conditioning, we confirmed appropriate targeting of our viral vectors to CA1 (**Fig. 3G**) and demonstrated that add-back of wildtype γCaMKII (*vs.* RFP) fully restores γCaMKIIQ285dop levels in dHPC to that of wildtype levels (**Fig. 3H**). We additionally explored whether such rescue may also be sufficient to restore deficits in γCaMKII trafficking to the synapse, which was previously observed in γCaMKIIQ285A mice. Our results indicated that this rescue approach was sufficient to restore γCaMKII synaptic trafficking to that of wildtype levels (**Fig. 3I**). Finally, we assessed whether restoring γCaMKIIQ285dop levels in CA1 may be sufficient to rescue contextual memory deficits observed in mutant animals. Following context re-exposure 24 hr post-training, mutant mice transduced with Lenti-γCaMKII were observed to display freezing levels similar to that of wildtype mice (transduced with Lenti-RFP), significantly reversing the memory deficits observed in γCaMKIIQ285A expressing Lenti-RFP alone (**Fig. 3J**).

## DISCUSSION

We feel that the data presented here provide a compelling case for protein dopaminylation as an important, yet previously undescribed, synaptic signaling mechanism in brain, with DA not only acting through cell surface-bound receptors but also infiltrating post-synaptic cells to directly modify proteins therein. We found that attenuation of γCaMKIIQ285dop – which is near exclusively synaptic – in dHPC results in disruptions in γCaMKII’s synaptic-to-nuclear signaling and its influence on neural transcriptional patterning, neuronal excitability, and context-dependent learning and memory. Based upon these data, we now hypothesize that upon dopamine uptake into cells and Ca^2+^-dependent cellular activation, TGM2 dopaminylates γCaMKII at position Q285 to promote γCaMKII trafficking to the membrane where it can then engage with βCaMKII. Reaching this previously described cellular ‘checkpoint’ is then critical for the proper downstream signaling of γCaMKII. Importantly, restoration of γCaMKIIQ285dop specifically within CA1 of mutant mice lacking the PTM rescues the expression of contextual fear memory in whole-body knock-in mice. We believe that this regional specificity helps to bolster the argument that γCaMKIIQ285dop functions as an acute mediator of learning and memory formation, and lends evidence to support the assumption that these effects are indeed dopamine-dependent, yet in a non-canonical, DA receptor-independent manner. Given that VTA-to-dHPC-dependent DA signaling has previously been shown to contribute importantly to contextual learning (*34*), it stands to reason that some of these effects may be mediated by dopaminylation of γCaMKII in dHPC. While compelling in and of itself, we wish to emphasize that similar dopaminylation-based regulation of the synaptic proteome is likely generalizable to many protein substrates beyond that of just γCaMKII, with these additional dopaminylated substrates (for which modified sites may exist both intra- and extracellularly) likely also contributing to important aspects of protein signaling and/or downstream gene expression, physiology, and behavior. For example, the dopamine transporter (DAT; Slc6a3) – a membrane-spanning protein that is critical for DA reuptake from the synapse to the cytosol of dopaminergic neurons – was identified as a substrate of dopaminylation in NAc (likely modified at VTA-to-NAc projection terminals). While the specific resides that are modified within DAT have yet to be elucidated, the possibility that dopaminylation of this transporter may contribute to its reuptake mechanism and, in turn, cellular physiology and behavior, is certainly intriguing. Another example is that of TGM2, which was found to be a substrate of dopaminylation in VTA, NAc and dHPC. Although much remains to be learned regarding how TGM2’s enzymatic activity and/or sub-cellular distribution are controlled within cells, such ‘automonoaminylation’ is perhaps suggestive of a potential monoamine sensing mechanism that may contribute to its regulation of monoaminylation states during periods of dynamic monoamine fluctuations in brain (and beyond). Finally, while our current data indicate a critical role for synaptic dopaminylation in adaptive brain plasticity (e.g., learning and memory), they may also implicate these phenomena in pathologies associated with altered monoaminergic signaling, although future studies will be necessary to fully elucidate these possibilities.

Importantly, this work does present with a few challenges/outstanding questions that will need to be addressed in follow up studies. First, while our *in vitro* and *in cellulo* validations of Bio-CO specificity towards dopaminylated *vs.* noradrenylated substrates indicated that the probe does not efficiently label noradrenylated proteins under the reaction conditions used in this study, it remains possible that noradrenylated substrates in brain (which have yet to be characterized owing to a lack of available tools/reagents for their investigation) are more robustly modified by TGM2 *vs.* that observed following NE treatments in primary neuronal culture. As such, future investigations will be needed to develop new tools that can fully distinguish between endogenously dopaminylated *vs.* noradrenylated protein *in vivo* Second, we acknowledge that our mutational approach perhaps represents an imperfect solution for testing the necessity of γCaMKIIQ285dop under normal physiological conditions, as the Q285A mutation might be expected to disrupt all possible monoaminylation events at this site (i.e., possibly disrupting serotonylation, histaminylation, etc.) if they are to occur *in vivo*. Unfortunately, however, the field currently lacks the tools necessary to *de novo* generate γCaMKIIQ285dop in brain/cells for causal analyses, although future studies implementing intein-based chemical methodologies (*41*), for example, may hold promise in this regard. In addition, while we found that the γCaMKIIQ285A mutation does not impact γCaMKII expression (RNA or protein) in mice or its intrinsic kinase activity *in vitro*, it remains possible that the mutation may affect other aspects of γCaMKII function *in vivo* that are not fully captured in our analyses and could be independent of its ability to be dopaminylated. Future studies may be needed to account for these additional possibilities. Finally, classical pathways for DA uptake via canonical transporter mechanisms (e.g., via SLC6A3/DAT) are believed to be restricted to certain populations of glial cells and the terminals of dopaminergic neurons, leaving little room for their utility in post-synaptic DA uptake, as suggested here (*42*). However, it is important to note that a myriad of non-canonical, high-capacity/low affinity monoamine transporters (e.g., organic cation transporters 2 and 3/OCT2 and OCT3, plasma monoamine transporter/PMAT) have been shown to be broadly expressed in brain and may also contribute to DA uptake into non-dopaminergic cells (*43, 44*). Specifically relevant to protein monoaminylation, it was recently shown that OCT3 – which is enriched at both plasma and nuclear membranes in neural cells (*45*) – plays a direct and causal role in the transport of 5-HT into astrocytes to promote the enzymatic addition of serotonylation on histone H3 (*46*). These previous findings suggest OCT3 as a prime target for future studies related to synaptic protein dopaminylation. Investigating these transporters, as well as potential alternative mechanisms of DA internalization (e.g., receptor internalization), in future studies will be paramount to dissecting the precise kinetics of dopaminylation events in non-monoaminergic cells in brain and may even represent novel targets for future therapeutic interventions aimed at treating pathologies associated with altered monoaminergic signaling.

Undoubtedly, additional monoamines – such as 5-HT, NE and/or histamine – will also be capable of similar actions in brain, including at the synapse, with such non-canonical monoaminylation events possibly representing a fundamental shift in our understanding of monoaminergic signaling writ-large. This is not to argue that canonical means of monoamine signaling/neuromodulation are not fundamentally important to cellular physiology, as they clearly are, but rather that protein monoaminylation likely also contributes importantly to cellular signaling and will require future intensive investigation. As such, with the creation of future tools aimed at targeting and dissecting mechanistic roles for these protein monoaminylation PTMs *in vivo*, we strongly believe that gaining a more detailed understanding of these phenomena may help to illuminate numerous mechanisms of brain physiology and disease that remain unresolved.

## ACKNOWLEDGMENTS

We would like to thank Drs. Paul Kenny and Paul Slesinger (ISMMS) for their experimental suggestions regarding this work. This work was partially supported by grants from the National Institutes of Health: R01 DA056595 (I.M.), R01 MH116900 (I.M.), R01 FM086868 (T.W.M.), R01 CA259365 (T.W.M.), F31 DA055462 (A.F.S.), F99 NS125774 (S.L.F.) and R01 CA251425 (R.J.D.). I.M is also supported by funds from the Howard Hughes Medical Institute.

## CONTRIBUTIONS

I.M. and A.F.S. conceived of the project with input from T.W.M. & R.E.T.

I.M. and A.F.S. designed the experiments and interpreted the data.

A.F.S., S.L.F., R.D.C., E.B., B.C., L.A.F., R.F., S.C., R.M.B., G.D.S., C.P., H.M., E.B., S.G.M.,

S.J.R., & I.M. collected and/or analyzed the data.

R.E.T., T.W.M., P.J.C. & R.J.D. contributed to probe synthesis and related experimental design.

S.L.F. performed the sequencing-based bioinformatics.

A.F.S., S.L.F. & I.M. wrote the manuscript with input from all other authors.

## DATA AVAILABILITY

The snRNA-seq data generated in this study have been deposited in the National Center for Biotechnology Information Gene Expression Omnibus (GEO) database under accession number GSE277319. Mass spectrometry proteomics data have been deposited to the ProteomeXchange Consortium via the PRIDE partner repository with the dataset identified as PXD055995. We declare that the data supporting findings for this study are available within the article and Supplementary Materials. Related data and code are available from the corresponding author upon reasonable request. No restrictions on data availability apply.

## INCLUSION & ETHICS STATEMENT

All collaborators associated with this work have fulfilled the criteria for authorship required by the American Association for the Advancement of Science. To obtain authorship, their participation in this study was deemed to be essential for the design and implementation of the work presented. Roles and responsibilities were agreed upon among collaborators ahead or during the research.

## Supplementary Materials for

### MATERIALS AND METHODS

#### Synthesis of Bio-Co probe

##### Materials

^1^H spectra were acquired on a Bruker DRX-600 spectrometer at 600 MHz for 1H. Thin layer chromatography (TLC) was performed on silica coated aluminum sheets (thickness 200 μm) or alumina coated (thickness 200 μm) aluminum sheets supplied by Sorbent Technologies, and column chromatography was carried out on Teledyne ISCO combiflash equipped with a variable wavelength detector and a fraction collector using a RediSep Rf high performance silica flash columns by Teledyne ISCO. LCMS/HPLC analysis for purity determination and HRMS was conducted on an Agilent Technologies G1969A high-resolution API-TOF mass spectrometer attached to an Agilent Technologies 1200 HPLC system. Samples were ionized by electrospray ionization (ESI) in positive mode. Chromatography was performed on a 2.1 × 150 mm Zorbax 300SBC18 5-μm column with water containing 0.1% formic acid as solvent A and acetonitrile containing 0.1% formic acid as solvent B at a flow rate of 0.4 mL/min. The gradient program was as follows: 1% B (0−1 min), 1−99% B (1−4 min), and 99% B (4−8 min). The temperature of the column was held at 50 °C for the entire analysis. The purity of all the compounds was ≥95%. N-[(1R,8S,9s)-Bicyclo[6.1.0]non-4-yn-9-ylmethyloxycarbonyl]-1,8-diamino-3,6-dioxaoctane was purchased from Sigma-Aldrich (CAS #: 1263166-93-3) and EZ-link desthiobiotin was obtained from ThermoFisher (CAS #: 80750-24-9).

##### Experiment

((1R,8S,9s)-bicyclo[6.1.0]non-4-yn-9-yl)methyl(2-(2-(2-(6-(5-methyl-2-oxoimida-zolidin-4-yl)hexanamido)ethoxy)ethoxy)ethyl)carbamate: A solution of ((1R,8S,9s)-bicyclo[6.1.0]non-4-yn-9-yl)methyl (2-(2-(2-aminoethoxy)ethoxy)ethyl)carbamate (133 mg, 1.23 eq, 410 µmol), 2,5-dioxopyrrolidin-1-yl 6-(5-methyl-2-oxoimidazolidin-4-yl)hexanoate (103.7 mg, 1.0 eq, 333.1 µmol) and triethylamine (139 µL, 3.0 eq, 999.2 µmol) in a minimum amount of DMF (0.5 mL) was stirred at 25 °C for 3 hours. After 3 hours, the reaction mixture was extracted with ethyl acetate and water. The ethyl acetate layer was collected, dried with sodium sulfate and filtered to give a filtrate. The filtrate was mixed with silica gel and dried by rotovap vacuum. The dry solid mixture was purified (solid loading) by the normal phase (*Teledyne ISCO* 12 gram column cat# 69-2203-312; 100% DCM for 3 minutes to 5% MeOH in DCM for 10 minutes to 10% MeOH in DCM for 6 minutes) to afforded ((1R,8S,9s)-bicyclo[6.1.0]non-4-yn-9-yl)methyl(2-(2-(2-(6-(5-methyl-2-oxoimidazolidin-4-yl)hexanamido)ethoxy)ethoxy) ethyl)carbamate (yield: 65 mg; 37.5%). All fractions were monitored by TLC. After development, (10% MeOH in DCM), TLC plates were stained with potassium permanganate and heating. Some impurities and the desired product were stained and appeared as brown spots. These fractions containing compounds that stained on TLC were then defined by direct MS only (the desired product unable to be detected at wavelength 220 nm and 254 nm). TLC Rf: 0.5 (10% MeOH in DCM). ^1^H NMR (MeOD-*d_4_*, 600 MHz) δ 4.07-4.04 (m, 1H), 3.74-3.70 (m, 1H), 3.62-3.57 (m, 1H), 3.52 (broad s, 4H), 3.44 (t, *J* = 5.19 Hz, 4H), 3.26 (t, *J* = 5.42 Hz, 2H), 3.19 (m, 2H), 2.21-2.07 (m, 2H), 2.11 (t, *J* = 7.45 Hz, 2H), 1.94-1.90 (m, 1H), 1.82-1.71 (m, 1H), 1.57-1.56 (m, 2H), 1.55-1.51 (m, 3 H), 1.42-1.38 (m, 2H),1.36-1.32 (m, 1H), 1.30-1.24 (m, 3H),1.23-1.19 (m, 3H),1.00 (d, *J* = 6.46 Hz, 3H), 0.89-0.77 (m, 2H). LCMS (ESI^+^): m/z 521.3634 (M+H)^+^.

##### Animals

Wildtype adult male mice (7-week-old) and pregnant females (E15; used for primary neuron culture experiments) (C57BL/6J) were purchased from The Jackson Laboratory. Homozygous γCaMKIIQ285A mice (bred to homozygosity) *vs.* wildtype littermate controls were generated as described below. Adult mice were group housed (5 per cage) and pregnant females singly-housed on a 12-hour light/dark cycle (lights on from 7:00 A.M. to 7:00 P.M.) at constant temperature (23 °C) with *ad libitum* access to food and water. Adult male mice (7-10 weeks old) were used for all experiments outlined in this manuscript and were sacrificed by rapid decapitation, with the exception of those involving primary cultured neurons (see below). All animal protocols were approved by the IACUC at the Icahn School of Medicine at Mount Sinai (ISMMS).

##### Generation of γCaMKIIQ285A knock-in mutant mice

The yCaMKIIQ285A knock-in mice were generated at the Stem Cell Engineering Core and the Mouse Genetics Core at the Icahn School of Medicine at Mount Sinai using CRISPR/Cas editing, as described in **Figure S3D**. Briefly, a synthetic single guide RNA (sgRNA) (Synthego) and recombinant *S. pyogenes* Cas9 nuclease (IDT, 1081059) were preassembled as a ribonucleoprotein (RNP) and injected into C57BL/6J blastocysts, together with a 200-mer single-stranded oligodeoxynucleotide (ssODN) carrying the CA>GC to generate the Q285A mutation and an ectopic HhaI recognition site used for screening. Founder mice were screened by restriction fragment length polymorphism (RFLP) using primers external to the ssODN (F1 and R1) and HhaI digestion of the PCR product to identify mice that underwent homology directed repair (HDR) after CRISPR/Cas9-induced double strand break (**Figure S3E**). Mice 7, 12, 41, 24, 31 and 51 showed an RFLP pattern compatible with an HDR event. To further validate the successful HDR event, mice 12, 31, 41 and 51 were subjected to targeted next generation sequencing (NGS) using primers F2 and R2. The sequencing data were analyzed with CRISPResso tool (*47*), and the results revealed that mice 12 and 41 carried the expected CA>GC change, but also an unwanted GA deletion as result of a nonhomologous end-joining (NHEJ) (**Figure S3F**). To determine if the GA deletion was on the same allele as the CA>GC change, the F2/R2 amplicon was cloned into a sequencing vector (pCR Blunt II-TOPO), and the plasmids derived from single *E. coli* clones were subjected to Sanger sequencing. The results demonstrated that mouse 41 carried the CA>GC change and the GA deletion on different alleles. Mouse 41 was therefore used as the founder for the yCaMKIIQ285A mouse colony (**Figure S3G**). To maintain the colony and identify mice carrying the yCaMKIIQ285A allele, mice were screened using the Q285A_F and R2 primers (specific for the HDR allele), as provided in the Table below.

##### Mouse genotyping strategy

DNA for genotyping was isolated from tail/digit biopsies. Tissues biopsies were digested in 200 μl of DirectPCR Lysis Reagent (Tail) (Viagen Biotech Inc, CA, USA) and 0.2 mg of proteinase K for 12–15 h at 55 °C followed by 1 hour at 85 °C for proteinase K inactivation. The digested tissues were directly used for PCRs. Thirty-five cycles of PCR amplification were carried out with a hybridization temperature of 60 °C.

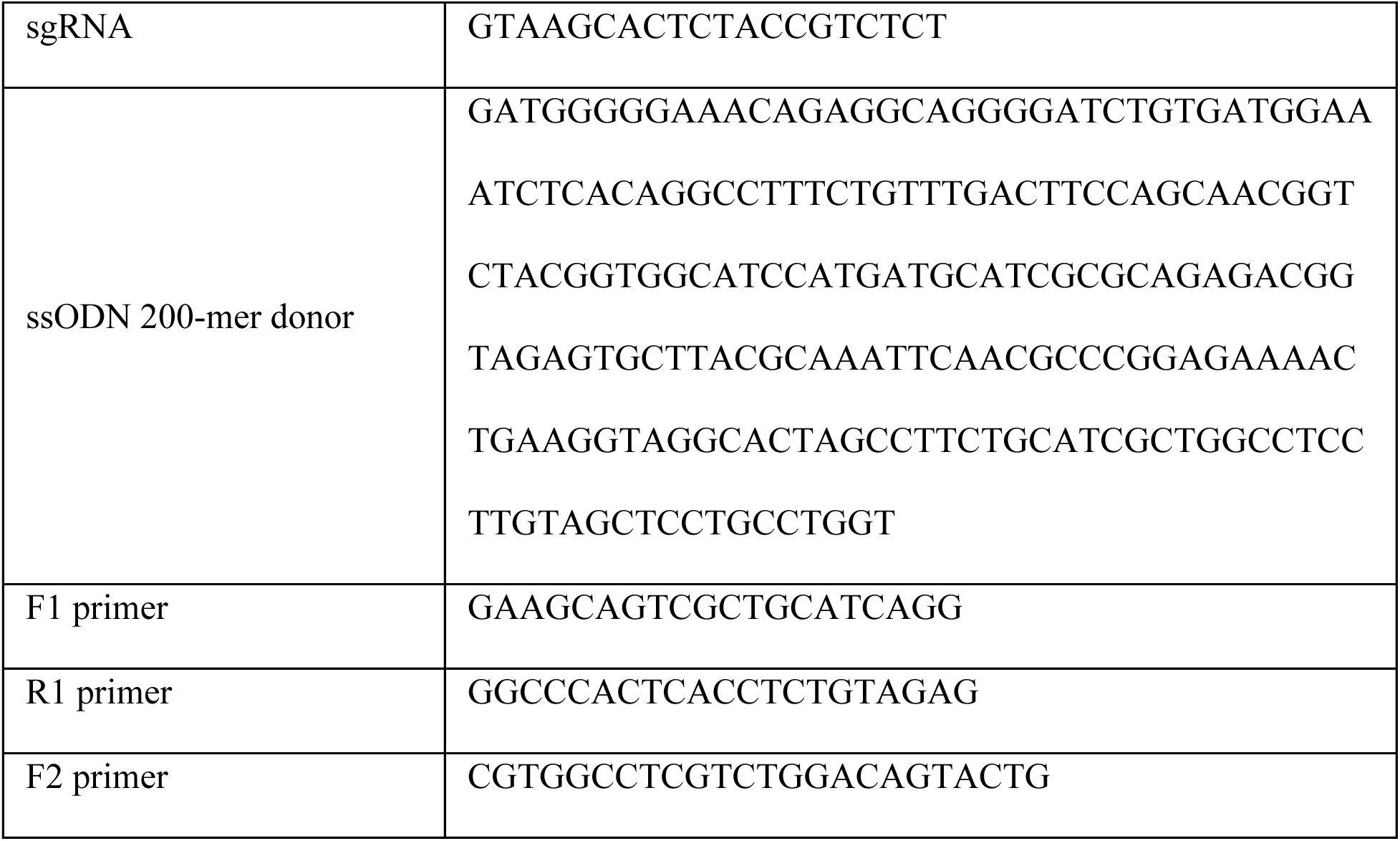
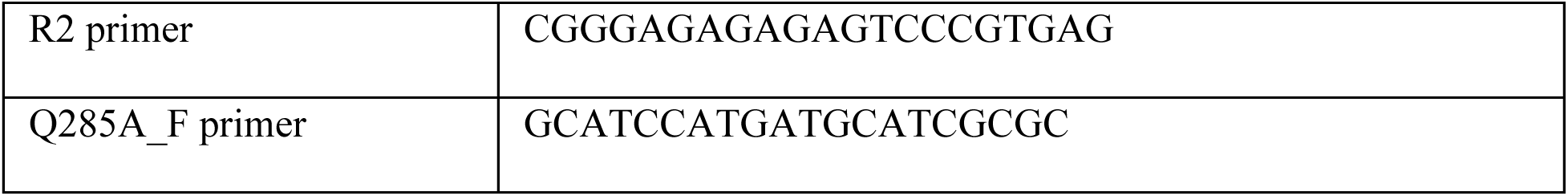

##### Primary mouse neuronal culture

E18 pregnant dams were anesthetized with isofluorane and then euthanized, followed by collection of embryos. Embryonic brains were extracted and microdessicted to isolate corticostriatal or hippocampal neurons. Cells were incubated in trypsin at 37 °C for 10 minutes, followed by trituration into a single-cell suspensions. Cells were then plated onto Poly-D-Lysine coated plates in DMEM+Glutamax (Gibco, 10566016), supplemented with penicillin/streptomycin (Gibco, 15140122) and 10% fetal bovine serum (Gibco, 26140079) at appropriate densities for given assays. Following 24 hours, media was aspirated and replaced with Neurobasal Plus Neuronal Culture System (Gibco, A3653401), supplemented with CultureOne (Gibco, A3320201) to suppress glial growth. Relevant experiments were performed on DIV14.

#### Protein purification

##### Purification of TGM2

The His6-tagged human TGM2 bacterial expression plasmid – pHis-hTG2 – was obtained from Addgene (Addgene plasmid # 100719) and was used to transform the *E. coli* strain BL21-Codon Plus (DE3)-RIPL (Agilent Technologies, 230280). Single colonies were picked from LB plates containing ampicillin, chloramphenicol and streptomycin, and were grown to early log phase (OD600 of 0.2-0.4) in LB medium in the presence of ampicillin at 37 °C. The temperature then was reduced to 16 °C and cultures were induced with 0.2 mM IPTG for overnight expression at 16 °C. Bacteria were harvested at 4800g for 20 minutes, and pellets were resuspended in 5 volumes of ice-cold lysis buffer containing 50mM Tris pH 7.5, 500mM NaCl, 50mM imidazole, 1mM PMSF and 1x cOmplete™ EDTA-free protease inhibitor cocktail (Sigma, 11836170001). Bacteria were lysed by sonication using a 5 second pulse, 5 second rest setting with 40% output, for a total of 20 minutes, on ice. Crude lysates were centrifuged at 25,000g for 30 minutes at 4 °C. The supernatant were passed through a syringe tip 0.45-micron filter unit and applied to a His-Trap affinity column (Cytiva, 17524701) equilibrated with lysis buffer (excluding protease inhibitors) on an Akta Purifier (Cytiva). Following washes of the column with lysis buffer, salt concentration were gradually reduced to 150 mM NaCl. Bound proteins were eluted with a gradient of Imidazole from 50 mM to 500 mM and eluates were collected on a fraction collector. Fractions containing the peak protein elution were pooled together and applied to a HiPrep 26/10 desalting column, equilibrated with storage buffer containing 50mM Tris pH 7.5 and 150mM NaCl. Eluate was concentrated using an Amicon Ultra centrifugal concentrator (Sigma) and applied to an ENrich SEC-650 10×300 Column (Biorad), equilibrated with storage buffer for separation of impurities based upon size exclusion. Peak TGM2 fractions were pooled together and concentrated again.

##### Purification of histone H3

Recombinant human histone H3.2 was expressed in *E. coli* BL21 (DE3), extracted by guanidine hydrochloride and purified by flash reverse chromatography, as previously described (*48*). The purified histones were analyzed by RP-LC-ESI-MS (*48*).

##### Purification of CaMKII

Human γCaMKII wildtype or Q285A, or αCaMKII wildtype, coding sequences were cloned into a mammalian expression vector (pcDNA3.1) under a CMV promoter, along with N-terminal FLAG-HA and C-terminal HA epitope tags (pIM.081 and pIM.082). Expi293F cells (ThermoFisher Scientific, A14527) were grown in Expi293 expression medium and were transiently transfected with the plasmid using Expifectamine transfection reagent, according to manufacturer’s protocol (ThermoFisher Scientific, A14524). Cells were harvested 48-72 hours post-transfection by centrifugation at 500g, washed once with D-PBS, and then the cell pellet was resuspended in ice-cold lysis buffer containing 50mM Tris pH 7.5, 300mM NaCl, 1% Triton X-100, 1mM DTT, cOmplete™ EDTA-free protease inhibitor cocktail (Sigma, 11836170001). Lysates were incubated on ice for 15 min before centrifugation at 25,000g for 30 minutes at 4 °C. The supernatant was passed through a syringe tip 0.45 micron filter unit and applied to ANTI-FLAG® M2 Affinity Gel (Millipore Sigma, A2220) equilibrated with lysis buffer, rotating at 4 °C for 2 hours. Following affinity binding incubation, lysate/bead mixtures were transferred to a chromatography column and washed extensively with lysis buffer (containing up to 0.5M NaCl), followed by washes with 50mM Tris pH 7.5, 150mM NaCl. Bead bound proteins were eluted with 0.1M Glycine, pH 3.5 solution. Eluate pH was immediately neutralized by adding 1/20th volume of 1M Tris, pH 8.0. Eluate was then passed through desalting columns equilibrated with the storage buffer consisting of 50mM Tris, pH 7.5, 150 NaCl and concentrated using Amicon Ultra centrifugal concentrator (Sigma).

##### *In vitro* TGM2 monoaminylation biochemical assays

TGM2 monoaminylation assays were performed by incubating 3µg of H3, αCaMKII, βCaMKII (ab132961) or γCaMKII with TGM2 (0.3µg) in 30µL of enzymatic buffer (25 mM Tris, 10 mM CaCl2, 10 mM DTT, 10 mM KCl, 5 mM MDC/5-HT/DA/NE/His, pH 7.8) for 1hr at RT. The samples were then prepped and run on a BisTris polyacrylamide gel and imaged using UV light (e.g., MDC assays) or western blotting for histone H3 (e.g., *in vitro* probe-IP experiment).

#### *In vitro* Kinase Assay

##### CaMKII Reaction Buffer

40mM Tris, pH 7.4 ATP 100µM

20mM MgCl2

.01mg/mL BSA 50µM DTT 100 µM CaCl_2_

Recombinant WT and Q285A h-γCaMKII reactions were performed using the ADP-Glo Kinase assay (Promega, V9101) according to the manufacturer’s instructions. Briefly, serial dilutions of rγCaMKII and rγCaMKIIQ285A (from 2.6µg to 16ng, 0ng) were plated in triplicate in CaMKII reaction buffer and incubated at RT for 30 minutes. Equivolume of ADP-Glo reagent was added and incubated at RT for 40 minutes. Kinase Detection Reagent were then added to each well and incubated for an additional 30 minutes at RT. Following incubation of kinase detection reagent, luminosity was measured using a Cytation3 (Biotek).

##### Luciferase Assay

Both WT and Q285A hippocampal primary neurons were treated with pGF-CREB-mCMV-EF1α-Puro lentiviral particles (Systems Biosciences, TR202va-p) on DIV2. At DIV 14, cells were treated with 500nM DA for 1 hour. Pierce Firefly Luciferase Kit (Thermo, 16175) was then used according to manufactures protocol. Luminescence was measured via a Cytation3 (Biotek).

##### Immunoprecipitation of dopaminylated proteins using the Bio-CO probe

Briefly, Dynabeads M280 streptavidin (Thermo #11206D) were aliquoted and washed 2x with LSB+5% bovine serum albumin (BSA), followed by resuspension of beads in 500 µL LSB+5% BSA. Bio-CO probe (or desthiobiotin for – probe conditions) at a concentration of 1mg/mL in 1:1 acetonitrile and water was added to the tube and rotated at 4 °C for 1 hour. While incubating, tissue samples (VTA, NAc, mPFC or dHPC; 2 mm punches) were homogenized in LSB + cOmplete EDTA-free protease inhibitor cocktail (Sigma #11873580001) via wand sonication. 5mg/mL NaIO_4_ was then added to all samples, and incubated for 10 minutes on ice, flicking halfway through. Following incubation of bead with probe/desthiobiotin, samples were washed 1x with LSB and resuspended in LSB. 1M DTT was then added to samples before the addition of the conjugated bead to sample. Samples were then rotated for 1 hour at 4 °C before being washed 6x with 1mL HSB. Following HSB washes, samples were washed with 500µL 0.2M glycine, pH 3.5, and the glycine was discarded. Beads were then prepped for downstream analysis (proteomics or western blotting).

**Immunoprecipitation of dopaminylated proteins using the Bio-CO probe**
|  |  |
| --- | --- |
| Low Salt Buffer (LSB) | 150 mM KCl<br>1% NP40<br>20mM HEPES pH 7.4<br>10 mM MgCl<sub>2</sub> |
| High Salt Buffer (HSB) | 500 mM KCl<br>1% NP40<br>20mM HEPES pH 7.4<br>10 mM MgCl<sub>2</sub> |

##### Bio-CO Probe Western Blotting and antibodies

Following Bio-CO probe IPs, beads were resuspended in TBS with LDS sample buffer (Invitrogen #NP0007) and sample reducing agent (Invitrogen #NP0004), and then heated at 98 °C for 7 minutes. Samples were placed back onto the magnet and supernatant loaded into 4-12% BisTris polyacrylamide gels (BioRad #3450125) for electrophoresis, followed by transfer to nitrocellulose membranes. Nitrocelluose membranes were then blocked in 5% BSA in TBS + 0.1% Tween 20 (TBST) for 1 hour followed by incubation with primary antibodies overnight at 4 °C. The following primary antibodies were used: mouse anti-Histone H3 (1:15,000, Abcam ab10799), anti-αCaMKII (1:1,000, CST 50049S, anti-βCaMKII (1:1,000, Abcam ab34703) or mouse anti-γCaMKII (1:5000, Abcam ab201966). The following day, membranes were washed in TBST 3x for 10 minutes and incubated for 1 hour at RT in TBST+5%BSA with anti-Mouse Alex Fluor™ 647 (1:10,000, Invitrogen A-21242), anti-Mouse Alex Fluor™ 546 (1:10,000, Invitrogen A-21123), and/or anti-Rabbit Alex Fluor™ 647 (1:10,000, Invitrogen A-21246). This was followed by 3 washes in TBST for 10 minutes each before quantification using fluorescence via a ChemiDoc MP imaging system (Biorad). Densitometry was used to quantify protein bands using Image J Software (NIH).

##### Mass spectrometry

Proteins on magnetic beads underwent reduction and alkylation, followed by partial on-bead digestion with Trypsin (Promega) for 3 hours at room temperature. Supernatant was extracted, and proteins were re-digested with Lys-C (Wako/Fuji) and Trypsin overnight. Digestion reactions were stopped with neat Trifluoroacetic acid. Samples were Solid Phase Extracted (*49*), prior to being analyzed by LC-MS/MS. A 50 min or 70 min analytical gradient (typical 2% B/98%A to 32%B/68%A, A: 0.1% formic acid, B: 80% acetonitrile/0.1% formic acid) was used and peptides were measured using high resolution/high mass accuracy mass spectrometers (Q-Exactive, Fusion Ascend or Fusion Lumos, ThermoFisher Scientific). Generated Data Dependent Acquisition data were searched and quantified using ProteomeDiscoverer (ThermoFisher Scientific)/Mascot (MatrixScience) or MaxQuant v.1.6 or higher. Depending on the experiments, UniProt’s mouse, rat or human databases were queried concatenated with Trypsin and Lys-C, as well as other potential contaminants (*50*). All quantitative data was generated as label free experiments. Analyses targeting specific peptides were designed as Parallel Reaction Monitoring (PRM) experiments and ion traces were extracted manually. Statistical analysis was carried out using Perseus v.2.0: prior to tests, signals were LOG2 transformed, possible contaminants were removed, and it was required that a protein must be measured in at least 2/3 of replicates in at least one sample group. Missing values were imputed.

##### Subcellular fractionations

Mouse dorsal hippocampus (dHPC) was homogenized in Buffer A (10mM HEPES (pH 7.9), 10 mM KCl, 1.5 mM MgCl_2_, 0.34 M sucrose, 10% glycerol, and 1mM EDTA, and 1X protease inhibitor cocktail) using a dounce. Following transfer to a microcentrifuge tube, Triton-100X was added to a final concentration of 0.1%, and the samples were incubated on ice for 30 minutes before being centrifuged for 5 min at 1300g at 4 °C. The supernatant was decanted and reserved on ice. Nuclear pellets were resuspended with Buffer A to wash, and then spun in the centrifuge for 5 min at 1300g at 4 °C, before being decanted and supernatant discarded. Nuclear pellets were then resuspended in LSB (see above) with protease inhibitor cocktail. For experiments involving synaptosomal fractions, supernatant was then centrifuged at 20000g at 4 °C for 10 minutes. Following centrifugation, supernatants were removed to a new tube (cytosolic fraction), and the crude synaptosomal pellet was resuspended in LSB. Crude synaptic and nuclear pellets were then sonicated to ensure homogeneity.

##### Western Blotting

Samples (10-20µg total protein) were loaded into BisTris polyacrylamide gels (Invitrogen or BioRad) for electrophoresis and then proteins were transferred to nitrocellulose membranes. Nitrocelluose membranes were then blocked in 5% BSA in TBS + 0.1% Tween 20 (TBST) for 1 hour followed by incubation with primary antibodies overnight at 4 °C. The following primary antibodies were used: mouse anti-Histone H3 (1:15,000, Abcam ab10799), mouse anti-CaMKII gamma (1:5000, ab201966), anti-PSD95 (1:500, Abcam ab18258; note that 5% nonfat dry milk in TBST was used for blocking and incubation of this antibody based on manufacture recommendations), rabbit anti-CaMKIIβ (1:1000 Abcam ab34703), chicken anti-GAPDH (1:10000Millipore AB2302), rabbit anti-pan-pCaMKIIT286 (1:1000 CST 12716S), Rabbit Anti-Sodium Potassium ATPase (1:10000 Abcam ab76020), rabbit anti-phospho-CREBS1333 (1:500 CST, 9198) or mouse anti-CREB (1:500 CST, 9104). The following day, membranes were washed in TBST 3 times for 10 minutes and incubated for 1 hour at RT in TBST+5%BSA with anti-Mouse Alex Fluor™ 546 (1:10,000, Invitrogen A-21123), anti-Rabbit Alex Fluor™ 647 (1:10000, A-21246) or anti-Chicken Alexafluor 488 (1:10000, Invitrogen A-11039). This was followed by 3 washes in TBST for 10 minutes each before quantification using fluorescence via a ChemiDoc MP imaging system (Biorad). Densitometry was used to quantify protein bands using Image J Software (NIH), and proteins were normalized to appropriate loading controls for fraction.

##### Co-immunoprecipitations

Dynabeads M280 Sheep-Anti Mouse IgG beads are washed with LSCB before being resuspended in LSCB+5% BSA with mouse anti-CaMKII gamma (Abcam ab201966) and rotated at 4 °C for 1 hour. Conjugated beads are then washed and resuspended in LSCB and added to mouse dHPC homogenized in LSCB. Sample-bead mixture incubated via rotation at 4 °C for 4 hour. Following incubation, supernatants were removed and discarded, and samples were washed with HSCB 6x. Samples are then resuspended in TBS and prepped for western blotting.

Co-immunoprecipitations
|  |  |
| --- | --- |
| Low Salt CoIP Buffer(LSCB) | 150 mM KCl<br>0.5% NP40<br>20mM HEPES pH 7.4<br>10 mM MgCl<sub>2</sub> |
| High Salt CoIP Buffer(HSCB) | 350 mM KCl<br>0.5% NP40<br>20mM HEPES pH 7.4<br>10 mM MgCl<sub>2</sub> |

#### Single-nuclei RNA-seq

##### Nuclei isolation and library preparation

Single-nuclei RNA-sequencing was performed on four 3mm dorsal hippocampal tissue punches from 8-week-old wildtype *vs.* γCaMKIIQ285A mutant mice. Nuclei were isolated using a modified version of the Matevossian and Akbarian sucrose density gradient isolation protocol (*51*). Briefly, tissues were thawed in 1 mL of lysis buffer (0.32 M Sucrose, 5 mM CaCl_2_, 3 mM magnesium acetate, 0.1 mM EDTA, 10 mM Tris-HCl pH 8, 1 mM DTT, 0.1% Triton X-100) with 50 μL of 25U/mL RNase inhibitor (Takara, cat#2313B) for 2 minutes in a 1 mL dounce homogenizer (Wheaton, cat# 357538). Tissues were gently homogenized with ∼20 strokes using a tight homogenizer. An additional 1 mL of lysis buffer was added to the homogenizer, another 10 strokes were performed and the 2 mL homogenates were transferred to 15 mL Open-Top Thinwall Polypropylene Tubes 16 x 95 mm (Beckman, cat# 361707). The homogenizers and douncers were washed with an additional 2 mL of lysis buffer, which was added to the Polypropylene Tube for a total of 4 mL of tissue homogenates. Homogenates were carefully underlaid with 9 mL of sucrose solution (1.8 M sucrose, 3 mM magnesium acetate, 1 mM DTT, 10 mM Tris-HCl, pH 8) and ultracentrifuged in a Sorvall™ WX+ at 24,000 rpm for 1 hr at 4 °C.

Following centrifugation, supernatants and the debris interphases were gently aspirated, resuspended in 1mL of resuspension buffer [0.02% bovine serum albumin in DPBS 25 µl of 25U/mL RNase inhibitor (Takara, cat#2313B)] and incubated on ice for 10 minutes. Resuspensions were filtered through a 35 µm nylon mesh filter (Corning, cat#352235) into a 1.5mL RNase/DNase-free microcentrifuge tube and centrifuged at 2,600 x g for 10 minutes at 4°C. Supernatants were removed and nuclear pellets were resuspended in 200 uL of resuspension buffer. 10 uL of nuclei were stained with Trypan Blue, and the quality and quantity of nuclei were assessed using a Countess 3 Automated Cell Counter.

Nuclear suspensions were then loaded onto a Chromium Single cell 3′ chip, version 3 (10X Genomics) and processed according to manufacturer’s protocol with a target of 8,000 cells. Single-nuclei libraries were prepared as per the 10X Chromium Next GEM Single Cell 3’ v3.1 (Dual Index) Protocol (CG000315 Rev A) being pooled onto a single 10B 100 Cycle Flowcell and sequenced using the Illumina NovaSeq X Plus system to obtain paired-end 2 × 100 bp reads.

### Data analysis

FastQ files were processed with the 10X Genomics Cell Ranger single-cell analysis pipeline (Cell Ranger v7.1.0) to demultiplex reads, align against the mouse genome (mm10-2020-A), deduplicate reads, and extract the gene expression matrix. Only confidently mapped, non-PCR duplicated reads with valid single-cell barcodes and UMIs were be used to generate gene-barcode matrices for further analysis. For each sample, we used the SoupX (*52*) (v1.6.2) package to automatically calculate the contamination fraction from the raw and filtered Cell Ranger output feature matrices, and then to remove background contamination from the count matrix. The Seurat pipeline (*53*) (v. 5.1.0) was used to carry out downstream analyses on the adjusted count matrices as Seurat objects. First, quality control metrics were calculated per sample, and nuclei were filtered based on the following conditions: nFeature_RNA > 250 & nCount_RNA > 1000 & nCount_RNA < 110000 & percent.mt < 5 & percent.ribo < 5. Individual sample counts were normalized with *NormalizeData*, top variable genes were identified with *FindVariableFeatures* (using the “vst” method), and genes were centered and scaled with *ScaleData,* followed by *RunPCA* dimensionality reduction and *RunUMAP.* Individual samples were then integrated with Harmony (*54*) (v1.2.0). Next, distinct cell types were identified using Louvain (*55*) clustering and visualized by UMAP (*56*). Nearest neighbors were identified, and clusters were identified by a shared nearest neighbor (SNN) modularity optimization based algorithm (*FindClusters).* Cell clusters were annotated as major hippocampal cell-types by expression of known cell-type markers and through Seurat’s reference mapping capability with a previously published hippocampal single-cell sequencing anchor dataset from the Allen Brain Map resource (*57*).

Astrocytes: Aldh1l1, Gfap, Fzd2

Microglia: C1qc

Oligodendrocytes: Myrf, Mag, Mog

OPCs: Pdgfra

Endothelial Cells: Flt4 (58)

Pericytes: Vtn (59)

Nueral progenitor Cells (60): Neurod1, Ccnd2

Gabaergic: Gad2

Neural Stem Cells (60): Id4, Sox2, Fgfr3

CA1(59): Wfs1, Ndst4

CA3 (57, 59): Iyd, Cpne4

Subiculum (57): Tshz2

For differential expression testing, we used Seurat’s pseudobulk capability with *AggregateExpression* to sum gene counts from all the cells from the same sample for each cell type cluster. We then used DESeq2 at the sample level to perform differential expression testing with multiple hypothesis testing adjusted using FDR<5%. The clusterProfiler package (v 2.1.6) was implemented for pathway analysis of identified DEGs. We used the fgsea package (*61*) (v 1.30.0) to run gene set enrichment analysis on the output from pseudobulk differential expression testing.

The hdWGCNA (*62*) package (v. 0.3.03) was utilized to identify and visualize co-expression gene network modules with a soft power threshold of 6 (threshold identified via *TestSoftPowers*), as well as to visualize module enrichment and expression of the modules across cell-types. The hdWGCNA package was also used to calculate the correlation between modules by annotated cell-type and the trait of mutation (γCaMKIIQ285A) by Spearman’s rank correlation coefficients, with Student’s *t*-tests and a *p* value of < 0.05 being considered statistically significant. Finally, we used the EnrichR function within hdWGCNA to perform pathway analysis on the genes within each module.

### Ex vivo whole-cell patch-clamp recordings

4-8 week old male transgenic (Q285A) and wild-type littermate mice were anesthetized using isoflurane. Brains were rapidly extracted, and coronal sections (250 microns) were prepared using a Compresstome (Precisionary Instruments Inc.) in cold (0-4 °C) sucrose-based artificial cerebrospinal fluid (SB-aCSF) containing 87 mM NaCl, 2.5 mM KCl, 1.25 mM NaH2PO4, 4 mM MgCl2, 23 mM NaHCO3, 75 mM Sucrose, 25 mM Glucose. After recovery for 60 minutes at 32°C in oxygenated (95% CO2 / 5% O2) aCSF containing 130 mM NaCl, 2.5 mM KCl, 1.2 mM NaH2PO4, 2.4 mM CaCl2, 1.2 mM MgCl2, 23 mM NaHCO3, 11 mM Glucose, slices were kept in the same medium at RT for the rest of the day and individually transferred to a recording chamber continuously perfused at 2-3 mL/min with oxygenated aCSF. Patch pipettes (5-7 MΩ) were pulled from thin wall borosilicate glass using a micropipette puller (Sutter Instruments) and filled with a K-Gluconate-based intra-pipette solution containing 116 mM KGlu, 20 mM HEPES, 0.5 mM EGTA, 6 mM KCl, 2 mM NaCl, 4 mM ATP, 0.3 mM GTP (pH 7.2). Cells were visualized using an upright microscope with an IR-DIC lens and illuminated with a white light source (Olympus for Scientifica) and using the MicroManager v2.0 software (https://micromanager.org/).

All recordings were made in dorsal CA1. Excitability was measured in current-clamp mode by injecting incremental steps of current (0–300 pA, +20 pA at each step). Whole-cell recordings were performed using a patch-clamp amplifier (Axoclamp 200B, Molecular Devices) connected to a Digidata 1550 LowNoise acquisition system (Molecular Devices). Signals were low pass filtered (Bessel, 2 kHz) and collected at 10 kHz using Axon pCLAMP 11 Software Suite (Molecular Devices). Electrophysiological recordings were extracted using Clampfit (Molecular Devices). All groups were counterbalanced by days of recording and all recordings were performed blind to experimental conditions.

### Contextual fear conditioning

WT and Q285A male mouse littermates were habituated to the testing chamber for 15 minutes. The following day, mice were trained over 5 conditioning trials, each consisting of pseudo-random interval, with a 2.0s, 0.9mA foot shock. Testing for conditioned fear response (freezing) occurred 24 hour following training. Mice were placed into the conditioned context for 5 minutes, and total time freezing was recorded (VFC chamber, Med Associates). Freezing is expressed as a percentage of the total test time.

### Viral transduction

28 days before contextual fear conditioning, training animals were anaesthetized via ketamine/xylazine (10/1 mg/kg) i.p., positioned in a stereotaxic frame (Kopf instruments), and 0.5µl of viral construct (lenti-RFP *vs.* lenti-γCaMKII-RFP) was infused bilaterally into dHPC using the following coordinates from bregma; anterior-posterior (AP) −2.2, medial-lateral (ML) ±1.3, dorsal-ventral (DV) −1.6. Mice received meloxicam (1 mg/kg) s.c. and topical antibiotic treatments for 3 days following surgery. Behavioral testing and transduced tissue collection occurred at least 28 days following transduction to ensure maximal infection.

### Immunohistochemistry/immunofluroscence

Mice were anesthetized with isoflurane and perfused with cold 1X PBS, followed by 4% PFA. The brains were post-fixed overnight in 4% PFA and then placed in a 30% sucrose/PBS solution for two days. After a 1x PBS wash, the brains were embedded in Tissue-Tek® O.C.T. Compound (Sakura, #4583) and sectioned at a thickness of 40 µm using a cryostat (Leica CM3050-S). Serial sections were collected from the dHP and stored at 4°C in 1X PBS with 0.01% sodium azide until immunofluorescence processing.

For immunofluorescence, brain sections were washed three times in 1X PBS for 10 minutes each at room temperature (RT). The sections were then permeabilized with 0.2% Triton X in PBS for 30 minutes, followed by a 1-hour incubation at RT in blocking buffer (0.3% Triton X, 3% normal donkey serum, 1X PBS). To amplify the fluorescent signal of the viral constructs, the sections were incubated overnight at 4°C with mouse anti-RFP primary antibodies (ThermoFisher, catalogue #MA5-15257, 1:500). The following day, after three washes in 1X PBS, the sections were incubated for 2 hours at RT on a slow shaker with secondary antibodies (donkey anti-mouse AlexaFluor680 – ThermoFisher A-21109; 1:1000) in blocking solution. Afterward, the sections were washed three times in 1X PBS for 10 minutes each and incubated with DAPI (1:10000, Thermo Scientific 62248) for 5 minutes. Finally, the sections were washed in 1X PBS and mounted on charged thermofrost slides using ProLong Gold Antifade Mountant (ThermoFisher, Cat. No. P36934).

Digital images were captured using a confocal microscope (Zeiss LSM 780, upright) with Zen Black software. Images of the dHP were acquired at 20x magnification using an air objective (Plan-Apochromat 20x/0.8), with 4×3 tiled images. Eight consecutive acquisitions were averaged for each image at a bit depth of 16 bits. The fluorescent signal in the representative images was visualized as follows: RFP with excitation (Ex) at 561 nm and emission (Em) at 632 nm, and DAPI with excitation (Ex) at 405 nm and emission (Em) at 498 nm. The images were saved in .tiff or. czi format. In the representative images, the scale bar is set to 500 µm.

### Statistics

Statistical analyses were performed using Prism GraphPad software. For experiments comparing only two conditions, two-tailed Student’s t tests were performed. For experiments involving more than two conditions, one-way or two-way ANOVAs were performed with subsequent *post hoc* analyses. Sequencing-based statistical analyses are described above. In biochemical, physiological, behavioral and snRNA-seq analyses, all animals used were included as separate *n*s (i.e., samples were not pooled). Significance was determined at p≤0.05. All bar/dot and line plot data are represented as mean ± SEM.

**Figure S1:**
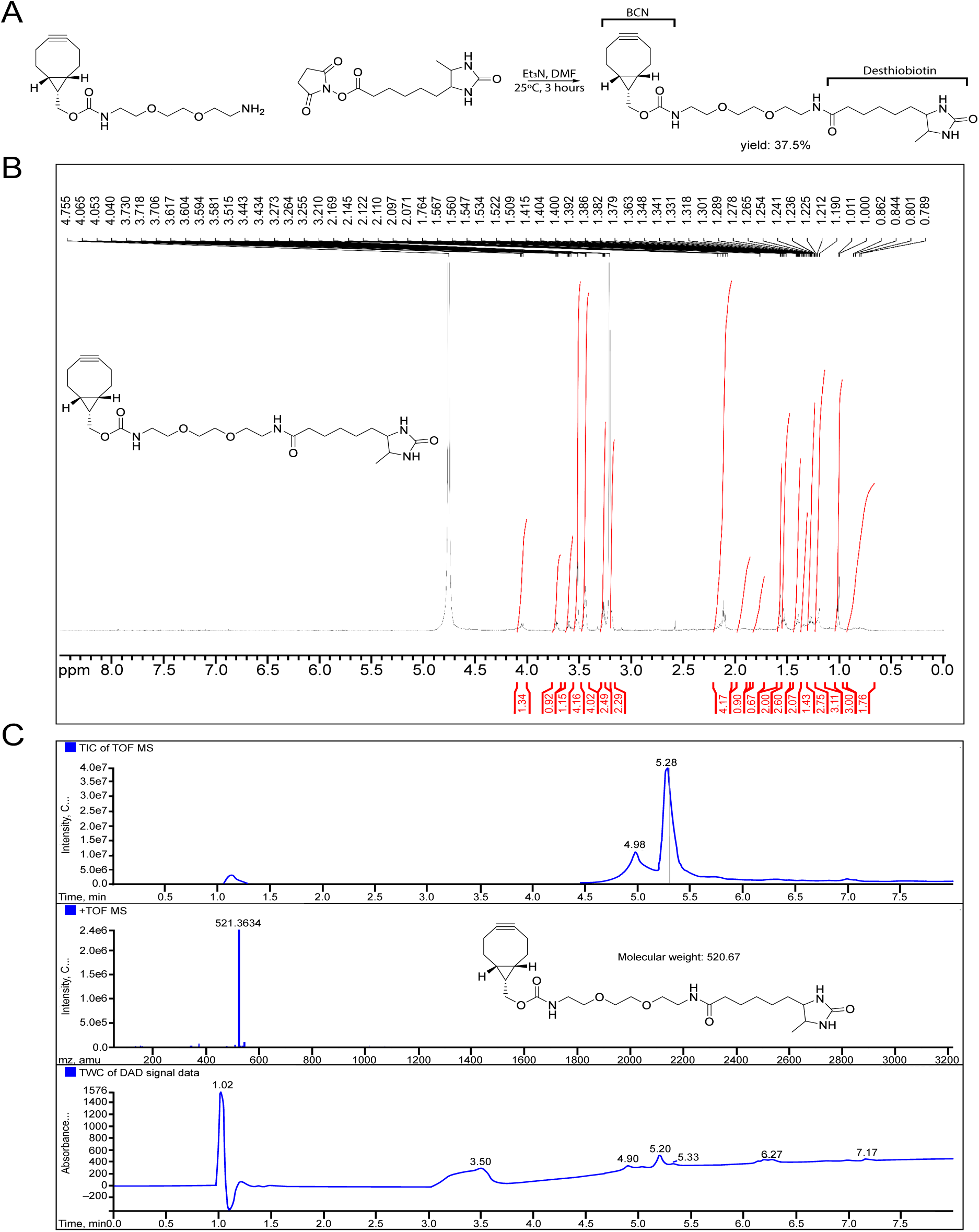
Synthesis and chemical validation of the Bio-CO probe. **A)** Chemical scheme for catecholaminylation probe synthesis: Bio-CO. **B)** ^1^H NMR spectroscopy validation of Bio-Co probe. **C)** LC-MS validation of Bio-CO probe.

**Figure S2:**
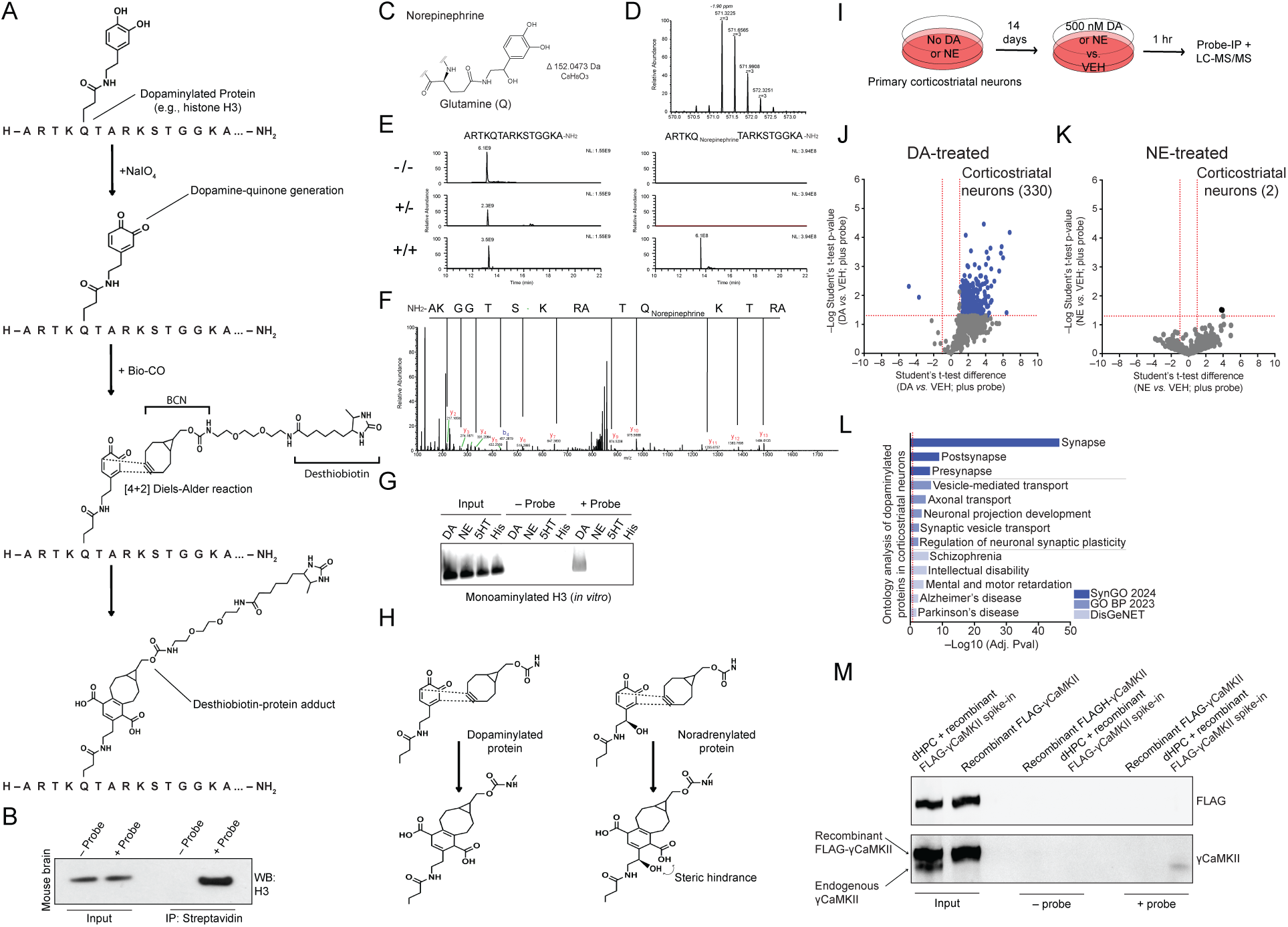
*in vitro, in cellulo* and *in vivo* validations of the Bio-CO catecholaminylation probe. **A)** Schematic of catecholaminylation probe-mediated bioorthogonal labeling of endogenous dopaminylated proteins. **B)** Catecholaminylation probe-mediated bioorthogonal labeling of endogenously dopaminylated histone H3 from mouse brain. Inputs = 1%; Streptavidin IPs were performed for – *vs.* + probe conditions, followed by blotting for H3. **C)** Proposed structure of noradrenylated glutamine. **D)** High resolution/high mass accuracy mass spectrum of the noradrenylated H3 tail peptide (ARTKQTARKSTGGKA-NH_3_; +NE/+TGM2 condition). **E)** Ion traces of the amidated H3 tail peptide with (right panels) and without (left panels) noradrenylated glutamine. Top, middle and bottom panels show signals measured under –/–, –/+ and +/+ (NE/TGM2) conditions, respectively. **F)** Tandem MS of the glutamine 5 noradrenylated H3 tail peptide (+NE/+TGM2 condition). Selected fragment ions (y and b) are labeled. Vertical lines within the peptide sequence are used to highlight simplified peptide fragmentation. **G)** Catecholaminylation probe-mediated bioorthogonal labeling of dopaminylated – but not serotonylated, histaminylated or noradrenylated – histone H3 *in vitro* following TGM2-dependent transglutamination. Inputs = 1%; Streptavidin IPs were performed for – *vs.* + probe conditions, followed by blotting for H3. **H)** Chemical rationale for diminished recognition of noradrenylated proteins by the catecholaminylation probe under the specific reaction conditions used in this study (e.g., steric hindrance). **I)** Schematic of monoamine (dopamine/DA or norepinephrine/NE) treatments of primary cultured mouse corticostriatal neurons, followed by probe IP-MS. **J)** Volcano plot of MS-identified proteins enriched by the catecholaminylation probe following treatments with DA *vs.* vehicle/VEH (Student’s t-test corrected for multiple comparisons; + probe: DA *vs.* VEH: *n* = 1%; *p*<0.05, fold-change >1). **K)** Volcano plot of MS-identified proteins enriched by the catecholaminylation probe following treatments with NE *vs.* vehicle/VEH (Student’s t-test corrected for multiple comparisons; + probe: NE *vs.* VEH: *n* = 1%; *p*<0.05, fold-change >1). **L)** Ontology analysis (SynGO 2024, GO BP 2023, DisGeNET) of MS-identified proteins enriched by the catecholaminylation probe following treatments with DA (FDR<0.05; Benjamini-Hochberg). **M)** Control experiment demonstrating that endogenous γCaMKII (dHPC), but not spike-in recombinant γCaMKII (purified using Expi293 cells, which lack endogenous monoaminylation owing to a lack of TGM2 expression) can be efficiency IP’d using the catecholaminylation probe. See **Figure S8** for uncropped blot images.

**Figure S3:**
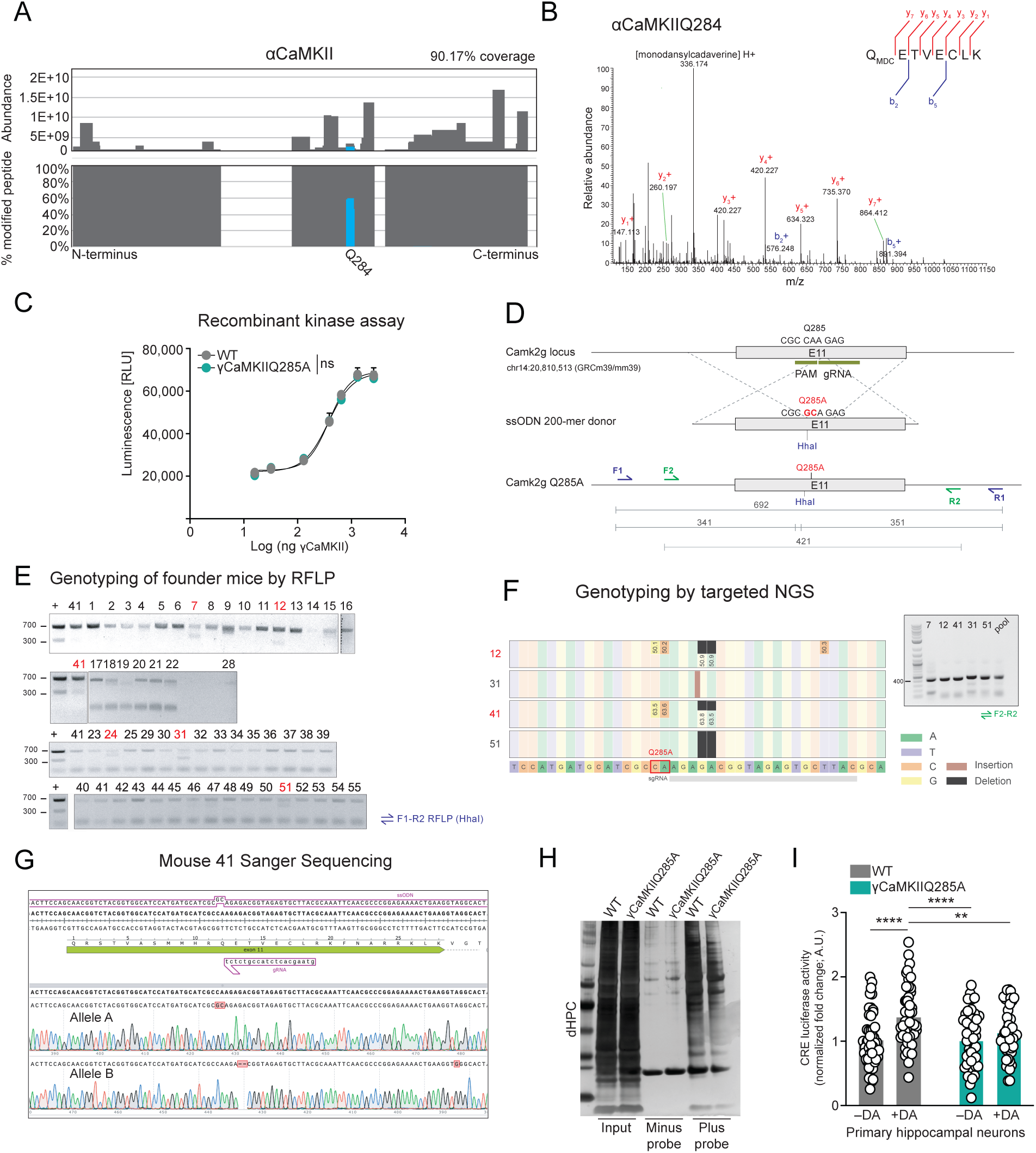
Generation of γCaMKIIQ285 knock-in mutant mice. **A)** Lego plot indicating the abundance and % of MDC modified peptide in αCaMKII (90.17% coverage) following TGM2-mediated transglutamination. Note that only αCaMKIIQ284 was observed to be modified by MDC. **B)** MS spectra for MDCylation of αCaMKII at glutamine 284. Y+ and b+ ions are annotated in red and blue, respectively. **C)** *in vitro* recombinant γCaMKII kinase activity assay, demonstrating that the Q285A mutation does not impact γCaMKII’s intrinsic kinase activity (*n* =3/γCaMKII concentration). The curves were not found to be statically different via nonlinear fit regression analysis. Data presented as mean ± SEM. **D)** Schematic of gene editing approach. Guide RNA and PAM sequences are indicated in green, the 200-mer ssODN donor used to mediate HDR contained the CA>GC change highlighted in red. The CA>GC generated the Q285A mutation, and the generation of HhaI recognition site was used for the screening. Primers used for the screening are indicated in purple and green. **E)** Genotyping of founder mice by restriction fragment length polymorphism (RFLP). Tail genomic DNA was amplified using F1 and R1 primers, followed by RFLP with HhaI. The expected bands (341 and 351 bp) for HDR events are detected in mice indicated in red. Positive control (+) is a spike-in of dsDNA containing the entire locus sequence spanning across F1 and R1 primers and containing the CA>GC change. **F)** Genotyping by targeted next generation sequencing. Mice 12, 31, 41 and 51 were selected for further validation. The amplicon generated using F2 and R2 primers of 421 bp was subjected to Illumina amplicon sequencing and analyzed using the CRISPResso tool. Mice 12 and 41 showed the expected HDR CA>GC change, but also a GA deletion in about half of the reads. **G)** Mouse 41 Sanger sequencing. The F2/R2 amplicon was cloned into a sequencing vector (pCR Blunt II-TOPO) and plasmids derived from single *E. coli* clones were subjected to Sanger sequencing, revealing that mouse 41 carried the CA>GC change and the GA deletion on different alleles. Mouse 41 was therefore used as the founder for the γCaMKIIQ285A mouse colony. **H)** Silver stain of catecholaminylation probe-mediated bioorthogonal labeling and enrichment of proteins from dHPC of WT *vs.* γCaMKIIQ285A mice, indicating that the Q285A mutation does not globally alter protein domainylation. Inputs =1%; Streptavidin IPs were performed for – *vs.* + probe conditions. **I)** CRE luciferase assay in WT *vs.* γCaMKIIQ285A primary cultured hippocampal neurons treated with dopamine *vs.* vehicle (VEH: WT *vs.* γCaMKIIQ285A, *n* = 47/genotype; DA: WT *vs.* γCaMKIIQ285A, *n* = 48 and 46/genotype, respectively). Analyzed by two-way ANOVA – main effects of treatment (F_1,184_ = 14.90, \*\*\**p* = 0.0002) and genotype (F_1,184_ = 3.886, \**p* = 0.0502), with significant post hoc analyses (uncorrected Fisher’s LSD) for: WT VEH *vs.* WT DA (\*\*\*\**p*<0.0001), Q285A VEH *vs.* WT DA (\*\*\*\**p*<0.0001) and WT DA *vs.* Q285A DA (\*\**p* = 0.0072). All line/bar plots presented as mean ± SEM. See **Figure S8** for uncropped blot images.

**Figure S4:**
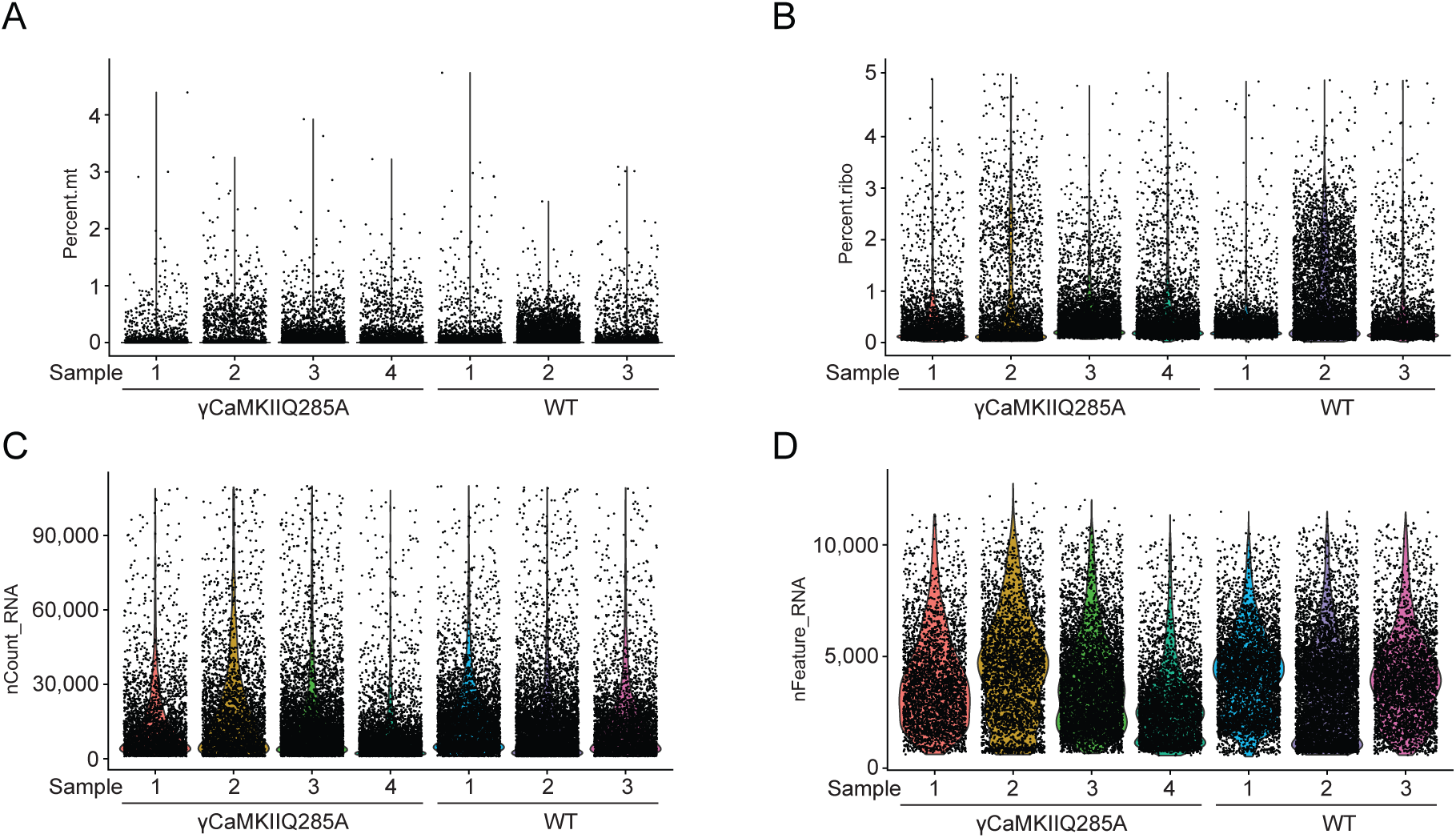
snRNA-seq quality control assessments. **A)** Violin plot showing the % of mitochondrial genes by sample (WT: *n* = 3 *vs.* γCaMKIIQ285A: *n* = 4) in snRNA-seq dataset after filtering. **B)** Violin plot showing the % of ribosomal RNA by sample (WT: *n* = 3 *vs.* γCaMKIIQ285A: *n* = 4) in snRNA-seq dataset after filtering. **C)** Violin plot showing the number of UMI by sample (WT: *n* = 3 *vs.* γCaMKIIQ285A: *n* = 4) in snRNA-seq dataset after filtering. **D)** Violin plot showing the number of detected genes by sample (WT: *n* = 3 *vs.* γCaMKIIQ285A: *n* = 4) in snRNA-seq dataset after filtering.

**Figure S5:**
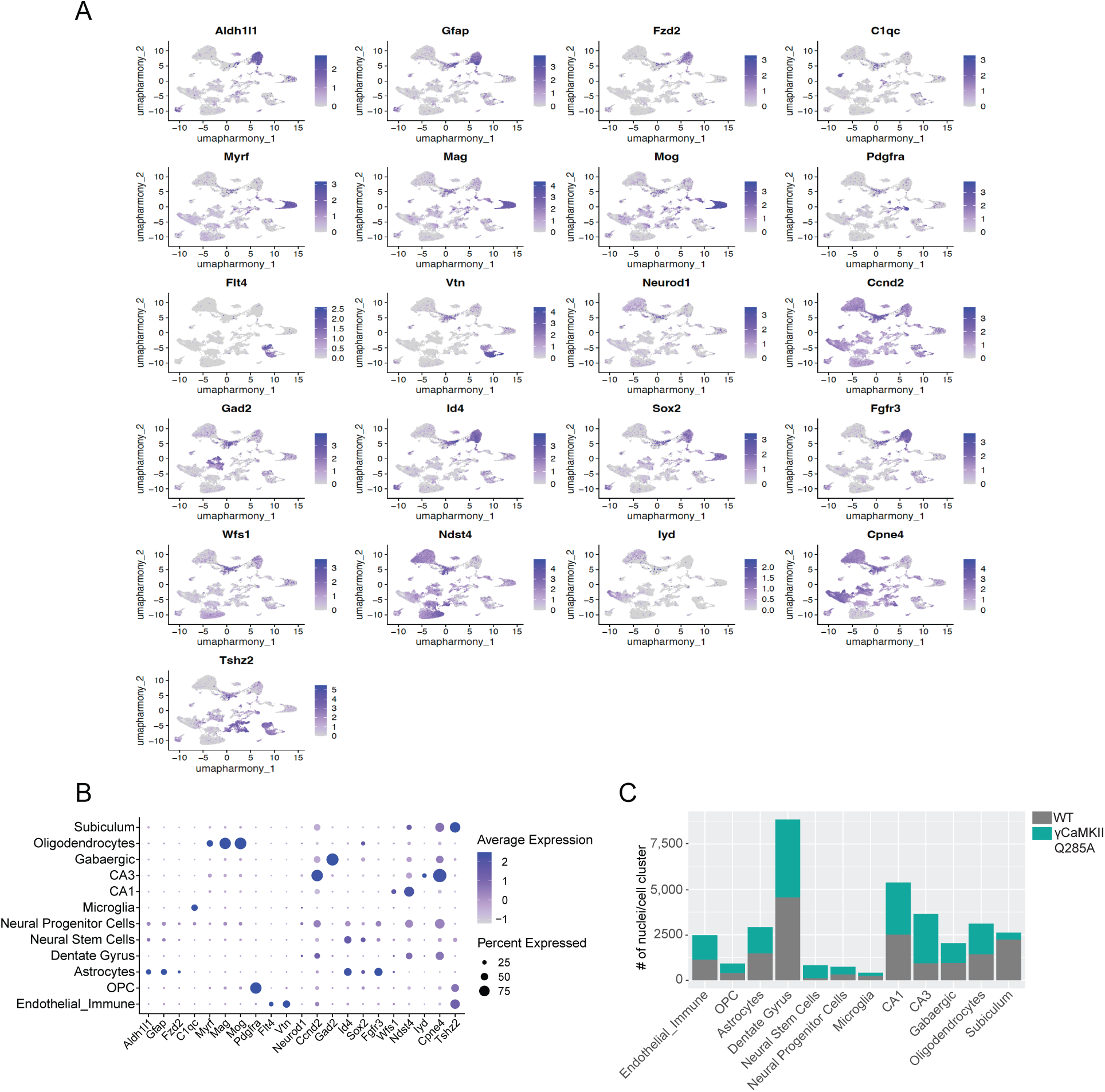
Cell cluster annotation from snRNA-seq dataset. **A)** Unsupervised clustering of snRNA-seq dataset (WT: *n* = 3 *vs.* γCaMKIIQ285A: *n* = 4) with cells colored by cell-type specific marker genes (see **Figure 2A** for cell cluster annotation by cell-type). **B)** Bubble plot indicating the average expression and % of cells expressing cell-type specific marker genes across annotated cell clusters (determined using unsupervised clustering). **C)** Stacked bar plot indicating the # of nuclei called per cell annotated cell cluster, separated by genotype (WT: *n* = 3 *vs.* γCaMKIIQ285A: *n* = 4).

**Figure S6:**
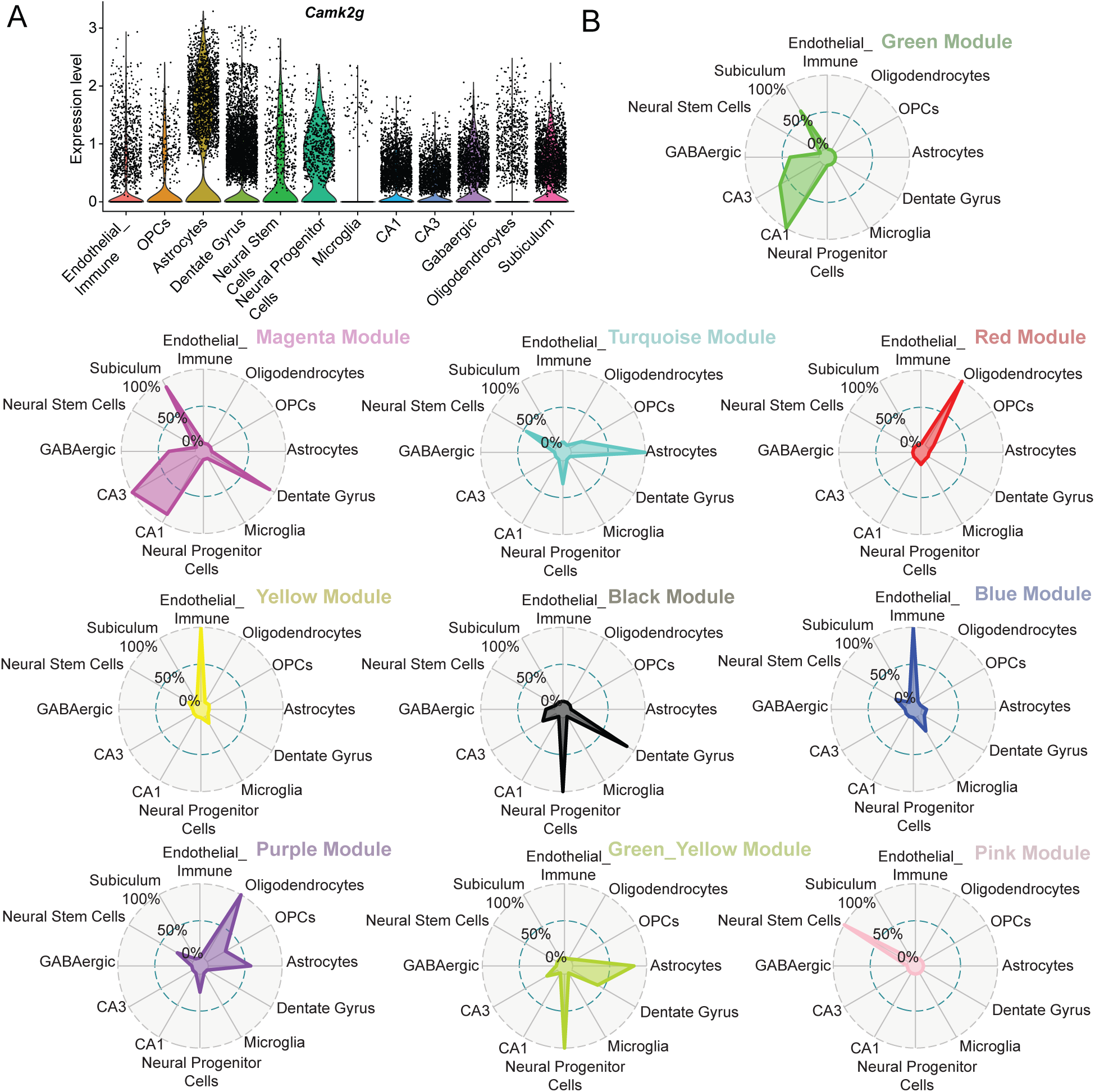
*γCaMKII* expression across cell clusters and WGCNA-identified module enrichment by cell cluster. **A)** *γCaMKII* RNA expression across annotated cell clusters in snRNA-seq dataset (WT: *n* = 3 *vs.* γCaMKIIQ285A: *n* = 4). Note that *γCaMKII* expression was not found to differ by genotype. **B)** Radar plots of WGCNA-identified module enrichment by annotated cell cluster (WT: *n* = 3 *vs.* γCaMKIIQ285A: *n* = 4). % enrichment of module genes by cell cluster is noted.

**Figure S7:**
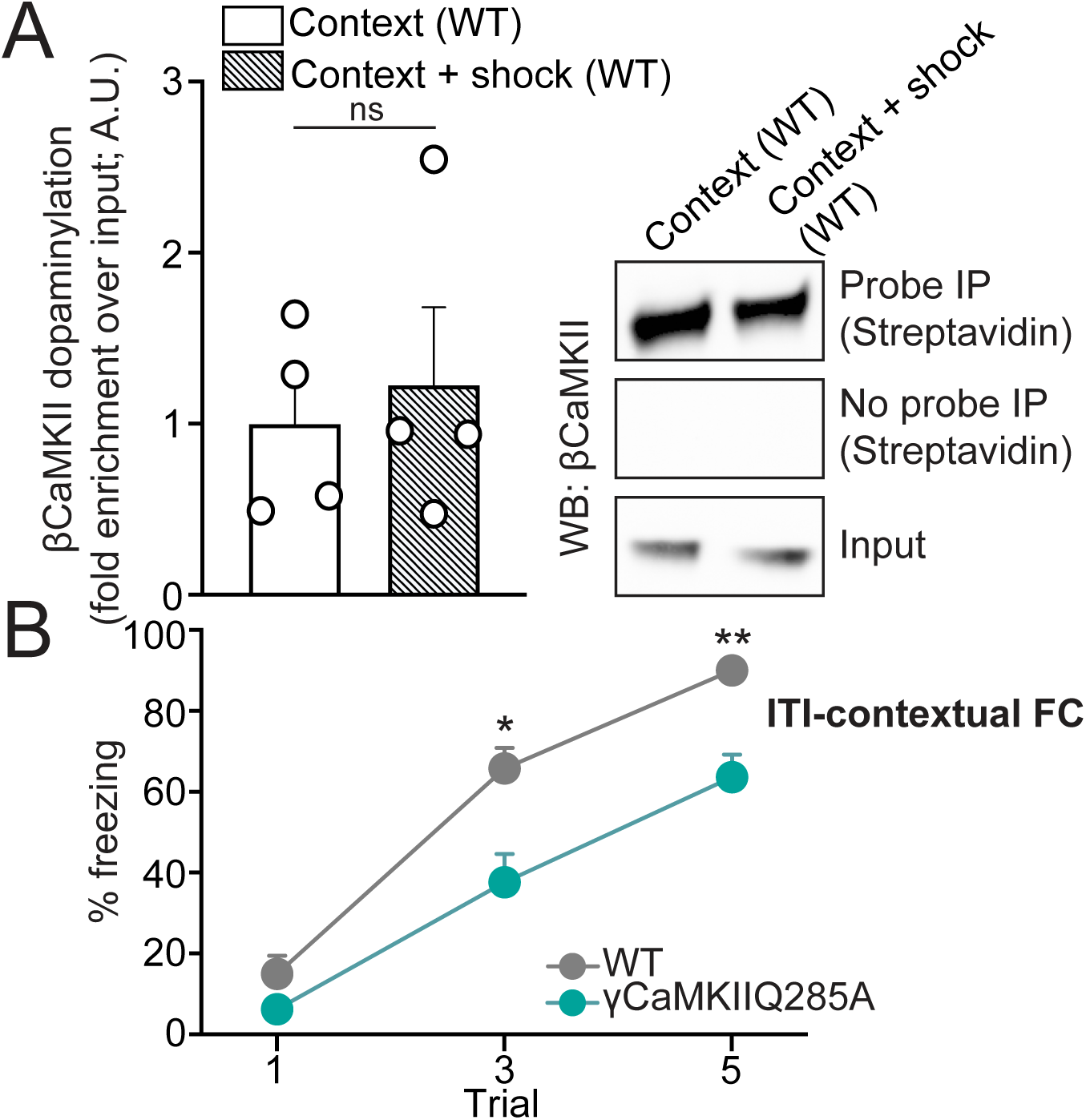
Contextual fear conditioning-induced dopaminylation is specific to γCaMKII and contributes to contextual learning in addition to contextual memory. **A)** Bio-CO probe-mediated bioorthogonal labeling of dopaminylated βCaMKII from dHPC of wildtype mice + (context + shock) *vs.* – (context alone) contextual fear conditioning training (1 hr post-training). Streptavidin IPs were performed for – *vs.* + probe conditions and βCaMKII levels were normalized to respective inputs (1%). *n* = 4/genotype – significance determined by unpaired Student’s t-test (t_6_ = 0.4300, *p* = 0.6822). **B)** Acquisition of contextual fear conditioning (average across inter-training interval/ITI trials) in wildtype *vs.* γCaMKIIQ285A mutant mice. Significance determined by repeated measures two-way ANOVA (main effect of trial x genotype; F_2,26_ = 4.302, \**p* = 0.0243), followed by post hoc analysis (Sidak’s MC test: \**p* = 0.0225, \*\**p* = 0.0076). All bar/line plots presented as mean ± SEM. See **Figure S8** for uncropped blot images.

**Figure S8:**
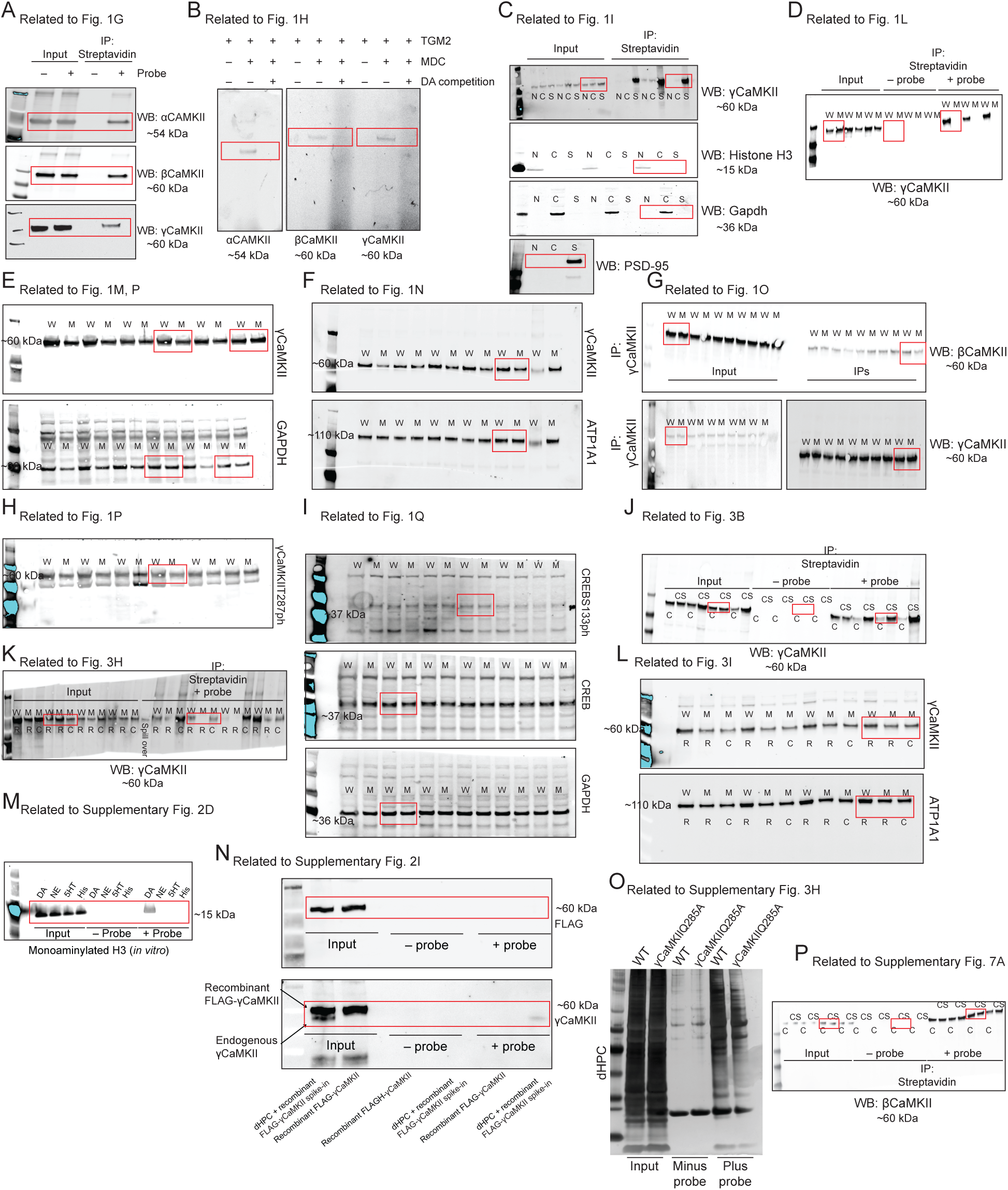
Uncropped blot images. **A)** Uncropped blots related to **Figure 1G**. **B)** Uncropped blots related to **Figure 1H**. **C)** Uncropped blots related to **Figure 1I**. **D)** Uncropped blots related to **Figure 1L. E)** Uncropped blots related to **Figure 1M,P**. **F)** Uncropped blots related to **Figure 1N**. **G)** Uncropped blots related to **Figure 1O**. **H)** Uncropped blots related to **Figure 1P**. **I)** Uncropped blots related to **Figure 1Q**. **J)** Uncropped blots related to **Figure 3B**. **K)** Uncropped blots related to **Figure 3H**. **L)** Uncropped blots related to **Figure 3I**. **M)** Uncropped blots related to **Supplementary** Figure 2D. **N)** Uncropped blots related to **Supplementary** Figure 2I. **O)** Uncropped blots related to **Supplementary** Figure 3H. **P)** Uncropped blots related to **Supplementary** Figure 7A. Red rectangles indicate the cropped regions of blots provided in the Main and Supplementary Figures.

### SUPPLEMENTARY DATA DESCRIPTIONS

**Data S1.** List of proteins identified by MS following Bio-CO labeling/enrichment in corticostriatal neurons plus DA treatment.

**Data S2.** List of proteins identified by MS following Bio-CO labeling/enrichment in corticostriatal neurons plus NE treatment.

**Data S3.** List of proteins identified by MS following Bio-CO labeling/enrichment in mouse VTA.

**Data S4.** List of proteins identified by MS following Bio-CO labeling/enrichment in mouse NAc.

**Data S5.** List of proteins identified by MS following Bio-CO labeling/enrichment in mouse mPFC.

**Data S6.** List of proteins identified by MS following Bio-CO labeling/enrichment in mouse dHPC.

**Data S7.** List of differentially expressed genes in dHPC comparing γCaMKIIQ285A *vs.* wildtype mice by annotated cell cluster (snRNA-seq pseudobulk analysis).

**Data S8.** List of genes within WGCNA modules identified in snRNA-seq dataset.

## REFERENCES AND NOTES

1. E. Dincer, M. Oguz, Y. Baris, Biochemical and pharmacological properties of biogenic amines. Biogenic Amines, (2018).

2. A. Al-Kachak, I. Maze, Post-translational modifications of histone proteins by monoamine neurotransmitters. Current Opinion in Chemical Biology 74, 102302 (2023).

3. G. D. Miller, Appetite regulation: hormones, peptides, and neurotransmitters and their role in obesity. American Journal of Lifestyle Medicine 13, 586–601 (2017).

4. J. S. Lin, C. Anaclet, O. A. Sergeeva, H. L. Haas, The waking brain: an update. Cell Mol Life Sci 68, 2499–2512 (2011).

5. M. Seyedabadi, G. Fakhfouri, V. Ramezani, S. E. Mehr, R. Rahimian, The role of serotonin in memory: interactions with neurotransmitters and downstream signaling. Experimental Brain Research 232, 723–738 (2014).

6. A. Gröger, R. Kolb, R. Schäfer, U. Klose, Dopamine reduction in the substantia nigra of parkinson’s disease patients confirmed by in vivo magnetic resonance spectroscopic imaging. PLOS ONE 9, e84081 (2014).

7. A. Sahu et al., The 5-Hydroxytryptamine signaling map: an overview of serotonin-serotonin receptor mediated signaling network. J Cell Commun Signal 12, 731–735 (2018).

8. I. V. Tabarean, Histamine receptor signaling in energy homeostasis. Neuropharmacology 106, 13–19 (2016).

9. M. O. Klein et al., Dopamine: functions, signaling, and association with neurological diseases. Cell Mol Neurobiol 39, 31–59 (2019).

10. K. W. Kroeze, K. Kristiansen, L. B. Roth, Molecular biology of serotonin receptors - structure and function at the molecular level. Current Topics in Medicinal Chemistry 2, 507–528 (2002).

11. L. A. Colgan, I. Putzier, E. S. Levitan, Activity-dependent vesicular monoamine transporter-mediated depletion of the nucleus supports somatic release by serotonin neurons. Journal of Neuroscience 29, 15878–15887 (2009).

12. A. B. Young, C. D. Pert, D. G. Brown, K. M. Taylor, S. H. Snyder, Nuclear localization of histamine in neonatal rat brain. Science 173, 247–249 (1971).

13. N. K. Sarkar, D. D. Clarke, H. Waelsch, An enzymically catalyzed incorporation of amines into proteins. Biochim Biophys Acta 25, 451–452 (1957).

14. M. J. Mycek, D. D. Clarke, A. Neidle, H. Waelsch, Amine incorporation into insulin as catalyzed by transglutaminase. Archives of Biochemistry and Biophysics 84, 528–540 (1959).

15. D. D. Clarke, M. J. Mycek, A. Neidle, H. Waelsch, The incorporation of amines into protein. Archives of Biochemistry and Biophysics 79, 338–354 (1959).

16. D. J. Walther et al., Serotonylation of small GTPases is a signal transduction pathway that triggers platelet alpha-granule release. Cell 115, 851–862 (2003).

17. R. Hummerich, J. O. Thumfart, P. Findeisen, D. Bartsch, P. Schloss, Transglutaminase-mediated transamidation of serotonin, dopamine and noradrenaline to fibronectin: evidence for a general mechanism of monoaminylation. FEBS Lett 586, 3421–3428 (2012).

18. T. S. Lai, C. S. Greenberg, Histaminylation of fibrinogen by tissue transglutaminase-2 (TGM-2): potential role in modulating inflammation. Amino Acids 45, 857–864 (2013).

19. L. A. Farrelly et al., Histone serotonylation is a permissive modification that enhances TFIID binding to H3K4me3. Nature 567, 535–539 (2019).

20. S. Zhao et al., Histone H3Q5 serotonylation stabilizes H3K4 methylation and potentiates its readout. Proc Natl Acad Sci U S A 118, (2021).

21. B. J. Lukasak et al., TGM2-mediated histone transglutamination is dictated by steric accessibility. Proc Natl Acad Sci U S A 119, e2208672119 (2022).

22. A. Al-Kachak et al., Histone serotonylation in dorsal raphe nucleus contributes to stress- and antidepressant-mediated gene expression and behavior. Nat Commun 15, 5042 (2024).

23. D. Sardar et al., Induction of astrocytic Slc22a3 regulates sensory processing through histone serotonylation. Science 380, eade0027 (2023).

24. H. C. Chen et al., Histone serotonylation regulates ependymoma tumorigenesis. Nature 632, 903–910 (2024).

25. A. E. Lepack et al., Dopaminylation of histone H3 in ventral tegmental area regulates cocaine seeking. Science 368, 197–201 (2020).

26. S. L. Fulton et al., Histone H3 dopaminylation in ventral tegmental area underlies heroin-induced transcriptional and behavioral plasticity in male rats. Neuropsychopharmacology 47, 1776–1783 (2022).

27. A. F. Stewart, A. E. Lepack, S. L. Fulton, P. Safovich, I. Maze, Histone H3 dopaminylation in nucleus accumbens, but not medial prefrontal cortex, contributes to cocaine-seeking following prolonged abstinence. Mol Cell Neurosci 125, 103824 (2023).

28. Q. Zheng et al., Histone monoaminylation dynamics are regulated by a single enzyme and promote neural rhythmicity. bioRxiv, 2022.2012.2006.519310 (2022).

29. A. Borrmann et al., Strain-promoted oxidation-controlled cyclooctyne–1,2-quinone cycloaddition (SPOCQ) for fast and activatable protein conjugation. Bioconjugate Chemistry 26, 257–261 (2015).

30. L. Burdine, T. G. Gillette, H. J. Lin, T. Kodadek, Periodate-triggered cross-linking of DOPA-containing peptide-protein complexes. J Am Chem Soc 126, 11442–11443 (2004).

31. N. Zhang et al., Bioorthogonal Labeling and Enrichment of Histone Monoaminylation Reveal Its Accumulation and Regulatory Function in Cancer Cell Chromatin. J Am Chem Soc, (2024).

32. N. Zhang et al., Chemical Proteomic Profiling of Protein Dopaminylation in Colorectal Cancer Cells. J Proteome Res 23, 2651–2660 (2024).

33. S. J. Russo, E. J. Nestler, The brain reward circuitry in mood disorders. Nat Rev Neurosci 14, 609–625 (2013).

34. F. J. P. Sayegh et al., Ventral tegmental area dopamine projections to the hippocampus trigger long-term potentiation and contextual learning. Nat Commun 15, 4100 (2024).

35. D. J. de Quervain, A. Papassotiropoulos, Identification of a genetic cluster influencing memory performance and hippocampal activity in humans. Proc Natl Acad Sci U S A 103, 4270–4274 (2006).

36. I. Voineagu et al., Transcriptomic analysis of autistic brain reveals convergent molecular pathology. Nature 474, 380–384 (2011).

37. M. Proietti Onori et al., The intellectual disability-associated CAMK2G p.Arg292Pro mutation acts as a pathogenic gain-of-function. Human Mutation 39, 2008–2024 (2018).

38. S. M. Cohen et al., Calmodulin shuttling mediates cytonuclear signaling to trigger experience-dependent transcription and memory. Nature Communications 9, 2451 (2018).

39. H. Ma et al., γCaMKII Shuttles Ca2+/CaM to the Nucleus to Trigger CREB Phosphorylation and Gene Expression. Cell 159, 281–294 (2014).

40. Z. Yao et al., A taxonomy of transcriptomic cell types across the isocortex and hippocampal formation. Cell 184, 3222–3241 e3226 (2021).

41. Y. David, M. Vila-Perello, S. Verma, T. W. Muir, Chemical tagging and customizing of cellular chromatin states using ultrafast trans-splicing inteins. Nat Chem 7, 394–402 (2015).

42. M. Jaber, S. Jones, B. Giros, M. G. Caron, The dopamine transporter: A crucial component regulating dopamine transmission. Movement Disorders 12, 629–633 (1997).

43. H. Bönisch, in Organic Cation Transporters in the Central Nervous System, L. C. Daws, Ed. (Springer International Publishing, Cham, 2021), pp. 119–167.

44. P. J. Gasser, Organic Cation Transporters in Brain Catecholamine Homeostasis. Handb Exp Pharmacol 266, 187–197 (2021).

45. P. J. Gasser, M. M. Hurley, J. Chan, V. M. Pickel, Organic cation transporter 3 (OCT3) is localized to intracellular and surface membranes in select glial and neuronal cells within the basolateral amygdaloid complex of both rats and mice. Brain Struct Funct 222, 1913–1928 (2017).

46. D. Sardar et al., Induction of astrocytic Slc22a3 regulates sensory processing through histone serotonylation. Science 380, eade0027 (2023).

47. K. Clement et al., CRISPResso2 provides accurate and rapid genome editing sequence analysis. Nat Biotechnol 37, 224–226 (2019).

48. Q. Zheng et al., Reversible histone glycation is associated with disease-related changes in chromatin architecture. Nat Commun 10, 1289 (2019).

49. J. Rappsilber, Y. Ishihama, M. Mann, Stop and go extraction tips for matrix-assisted laser desorption/ionization, nanoelectrospray, and LC/MS sample pretreatment in proteomics. Anal Chem 75, 663–670 (2003).

50. J. Bunkenborg, G. E. Garcia, M. I. Paz, J. S. Andersen, H. Molina, The minotaur proteome: avoiding cross-species identifications deriving from bovine serum in cell culture models. Proteomics 10, 3040–3044 (2010).

51. A. Matevossian, S. Akbarian, Neuronal nuclei isolation from human postmortem brain tissue. Journal of visualized experiments: JoVE, (2008).

52. M. D. Young, S. Behjati, SoupX removes ambient RNA contamination from droplet-based single-cell RNA sequencing data. Gigascience 9, giaa151 (2020).

53. T. Stuart et al., Comprehensive Integration of Single-Cell Data. Cell 177, 1888–1902.e1821 (2019).

54. I. Korsunsky et al., Fast, sensitive and accurate integration of single-cell data with Harmony. Nature methods 16, 1289–1296 (2019).

55. V. D. Blondel, J.-L. Guillaume, R. Lambiotte, E. Lefebvre, Fast unfolding of communities in large networks. Journal of Statistical Mechanics: Theory and Experiment 2008, P10008 (2008).

56. E. Becht et al., Dimensionality reduction for visualizing single-cell data using UMAP. Nature Biotechnology 37, 38–44 (2019).

57. Z. Yao et al., A taxonomy of transcriptomic cell types across the isocortex and hippocampal formation. Cell 184, 3222–3241. e3226 (2021).

58. M. Xiang et al., A Single-Cell Transcriptional Roadmap of the Mouse and Human Lymph Node Lymphatic Vasculature. Front Cardiovasc Med 7, 52 (2020).

59. E. Cid et al., Sublayer-and cell-type-specific neurodegenerative transcriptional trajectories in hippocampal sclerosis. Cell reports 35, (2021).

60. B. Artegiani et al., A single-cell RNA sequencing study reveals cellular and molecular dynamics of the hippocampal neurogenic niche. Cell reports 21, 3271–3284 (2017).

61. G. Korotkevich et al., Fast gene set enrichment analysis. biorxiv, 060012 (2016).

62. S. Morabito, F. Reese, N. Rahimzadeh, E. Miyoshi, V. Swarup, hdWGCNA identifies co-expression networks in high-dimensional transcriptomics data. Cell reports methods 3, (2023).

